# Pax37 gene function in *Oikopleura dioica* supports a neuroepithelial-like origin for its house-making Fol territory

**DOI:** 10.1101/2023.07.18.549157

**Authors:** David Lagman, Anthony Leon, Nadia Cieminska, Wei Deng, Marios Chatzigeorgiou, Simon Henriet, Daniel Chourrout

**Author notes:** Current address: Department of Clinical Science, University of Bergen, Bergen, NO-5020, Norway.

## Abstract

Larvacean tunicates feature a spectacular innovation not seen in other animals - the trunk oikoplastic epithelium (OE). This epithelium produces a house, a large and complex extracellular structure used for filtering and concentrating food particles. Previously we have shown that several homeobox transcription factors may play a role in patterning the OE. Among these are two *Pax3/7* duplicates that we named *Pax37A* and *Pax37B*. The vertebrate homologs, *PAX3* and *PAX7*, are involved in developmental processes related to neural crest and muscles. In the ascidian tunicate *Ciona robusta*, *Pax3/7* has been given a role in development of cells deriving from the neural plate border including trunk epidermal sensory neurons and tail nerve cord neurons as well as in neural tube closure. Here we have investigated the roles of *Pax37A* and *Pax37B* in the development of the OE using CRISPR-Cas9, analyzing scRNA-seq data from wild-type animals that were compared with scRNA-seq data from *C. robusta*. We revealed that *Pax37B* but not *Pax37A* is essential for the differentiation of cell fields that produce the food concentrating filter of the house: the anterior Fol, giant Fol and Nasse cells. Lineage analysis supports that expression of *Pax37* is under influence of Wnt signaling and that Fol cells have a neuroepithelial-like transcriptional signature. We propose that the highly specialized secretory epithelial cells of the Fol region either maintained or evolved neuroepithelial features as do “glue” secreting collocytes of ascidians. Their development seems to be controlled by a GRN that also operates in some ascidian neurons.

## 1. Introduction

Tunicates, the sister group of all vertebrates ^1^, contain a wide range of species (∼2100) that is divided into three classes, Ascidians, Thaliaceans and Larvaceans ^1^. Larvaceans are pelagic and the only tunicates retaining a chordate body plan throughout life. Several times per day, they secrete a large gelatinous extracellular structure called “the house” from a highly organized epithelium called the “oikoplastic epithelium” (OE) that covers most of the trunk. The house, used for filtering and concentrating food particles from the seawater, is made of cellulose and a large set of specialized glycoproteins that were named oikosins ^2,3^. Oikosins have been described in the larvacean *Oikopleura dioica* and their sequences do not show similarities with other known proteins ^2^.

The OE morphology shows some variation in different larvacean families. It was thoroughly described in oikopleurids where it always exhibits the same distinct cellular fields. Two major fields of the OE are the Fol’s and the Eisen’s oikoplast. The Fol’s oikoplast produces the food concentrating filter (*fcf*), and the Eisen’s oikoplast produces the inlet filter (*if*) ^4^. Based on cell size and nuclear morphologies, both fields are divided further into several subfields. The Fol’s oikoplast comprises the giant Fol, Nasse cells and the posterior Fol (Fig. 1) ^5,6^. The nuclei of cells located in the OE are highly polyploid due to endoreduplication ^7^. How the OE gets established has been the object of recent studies in *Oikopleura dioica*. It begins to appear ∼4 hours after fertilization (hpf) and has already at 9-10 hpf the number of cells observed in the adult OE ^5,6^. In earlier studies, we identified several homeobox transcription factors that are transiently and specifically expressed in the developing OE, suggesting their involvement in a highly organized patterning ^5,8^. The first confirmations of such a role came from RNA interference experiments showing that the *Prop-a* and *Prop-b* genes are important for the normal development of the dorsal midline of the epithelium and the expression of a oikosin specific of this region ^5^.

**Figure 1:**
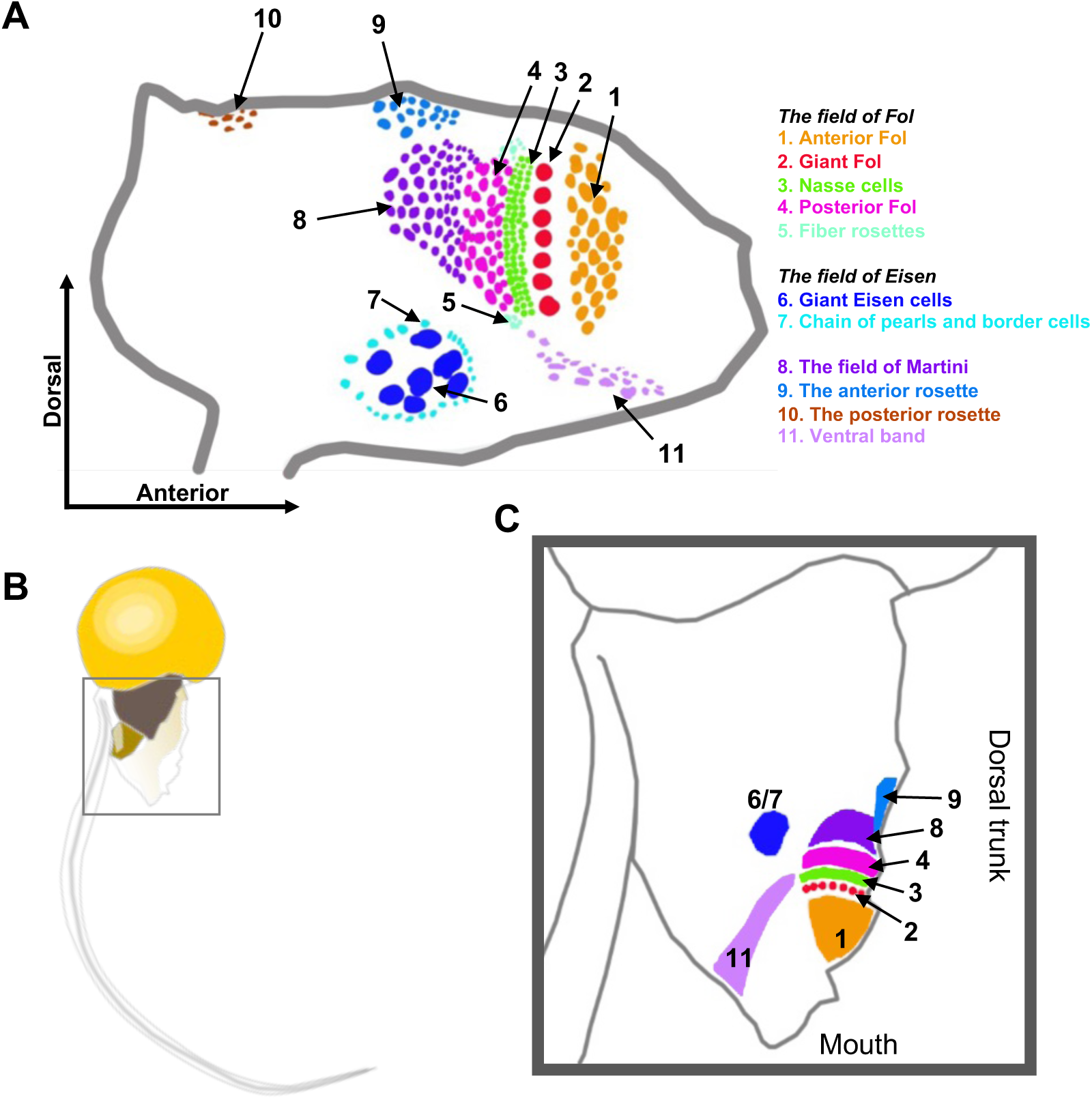
Schematic over the trunk oikoplastic epithelium of *O. dioica*. A) Detailed schematic of the epithelium with important oikoplastic fields labelled. Colored dots represent nuclei of cells in these regions. B) location of the trunk epithelium in the adult animal. C) Simplified schematic of the location of the largest fields in the adult trunk epithelium. Colors are the same as the names in the key.

Paired box proteins all have one or two DNA binding domains, either only a paired domain or a paired domain together with complete or partial homeodomain ^9^. The PAX3 and PAX7 proteins of vertebrates have a complete homeodomain as have their orthologs in tunicates and cephalochordates ^10,11^. The PAX3 and PAX7 of vertebrates play an important role in the induction of the neural crest and in the maintenance and differentiation of cells that migrate out of it to form the various neural crest derived tissues and muscles ^12^. The ascidian tunicate *Halocynthia roretzi* has a single *Pax37* gene, whose expression begins during early gastrula and continues until and during the larval stage ^13^. Expression is restricted to cell populations of the neural plate border that will develop into the dorsal neural tube, muscle cells and epidermal cells ^13^. *Pax37* is also expressed in dorsal epidermal cells that will give rise to epidermal sensory neurons ^13^. A recent study in *Ciona robusta* ^14^ showed an additional role in neural tube closure ^15^. *PAX3* expression in the neural plate border of vertebrates is induced by the canonical and non-canonical Wnt signaling pathways ^16^. This direct control of *Pax37* expression has not been reported for *PAX37* in ascidians, at least thus far. However, inhibiting Wnt signaling in *C. robusta* during gastrulation indicated a possible change of fate of the cells that later will develop into the anterior apical trunk epidermal sensory neurons (aATENs) ^17^. Since the development of these cells involved Pax37 ^18^, this could indicate a similar role of Wnt signaling.

The neural plate has not been identified in the developing *O. dioica* embryo ^19^ but data from later stages of development show that *Pax37A* and *Pax37B* are expressed in the dorsal trunk epiderm ^5,8^ from the hatching stage, as also observed in ascidians at the tadpole stages. Later during the larval stage, *Pax37A* and *Pax37B* are expressed in the epithelial fields that will develop into the different parts of Fol’s oikoplast ^5,8^.

Here we studied the roles of Pax37 genes in the development of the OE using CRISPR-Cas9 and single cell RNA-seq to investigate if: 1) these genes play a role in the development of Fol’s oikoplast and 2) have the cells of the Fol’s oikoplast evolved as new cell types as a result of the co-option of Pax3/7 and/or a gene regulatory network involving Pax3/7. We found that *Pax37B* but not *Pax37A* is essential for the differentiation of the giant Fol and Nasse cells and further survival of the animal. We also reveal that components of the canonical and non-canonical Wnt signaling pathways are among *Pax37B+* cell lineage markers. The transcriptomic profile of the *Pax37B+* epidermal clusters shares similarities with both epidermis and epidermal neurons in the *C. robusta* tadpole, indicating either a further specialization of these epidermal neuronal cells in larvaceans or a co-opted cascade of transcriptional regulation in an originally epidermal cell type.

## 2. Results

### 2.1 *Pax37* has experienced multiple duplications in larvacean genomes

Through blast searches in all available larvacean genomes and standard annotation procedures ^20,21^, we identified the presence of Pax37 gene duplicates, as reported earlier for *O. dioica*. Orthologs of *O. dioica Pax37A/B* type genes were found in all but two of the investigated oikopleurid genome assemblies. In the *Mesochordaeus erythrocephalus* and *Bathochordaeus sp*. only *Pax37B* was identified. A more variable expansion of additional *Pax37-like* genes was observed among oikopleurid genomes, leading to between two and eight genes (data not shown). In the available fritillarid genome assemblies (*Fritillaria borealis* and *Appendicularia sicula*) two Pax37 genes were identified that both appear to be equally closely related to the *Pax37A/B* genes. This indicates that the oikopleurid *Pax37* repertoire was established after the split from the fritillarids and that their duplication events are independent. Notably, as for many genes of larvaceans, the sequences of *O. dioica* predicted Pax37 proteins seem to have diverged rather rapidly from their orthologs in other chordate groups. However, those of *Pax37A/B* type appear much less divergent than those of *Pax37C/D*. For a phylogenetic analysis of the *O. dioica*, giant larvacean and fritillarid Pax37 sequences see Fig. 2A.

**Figure 2:**
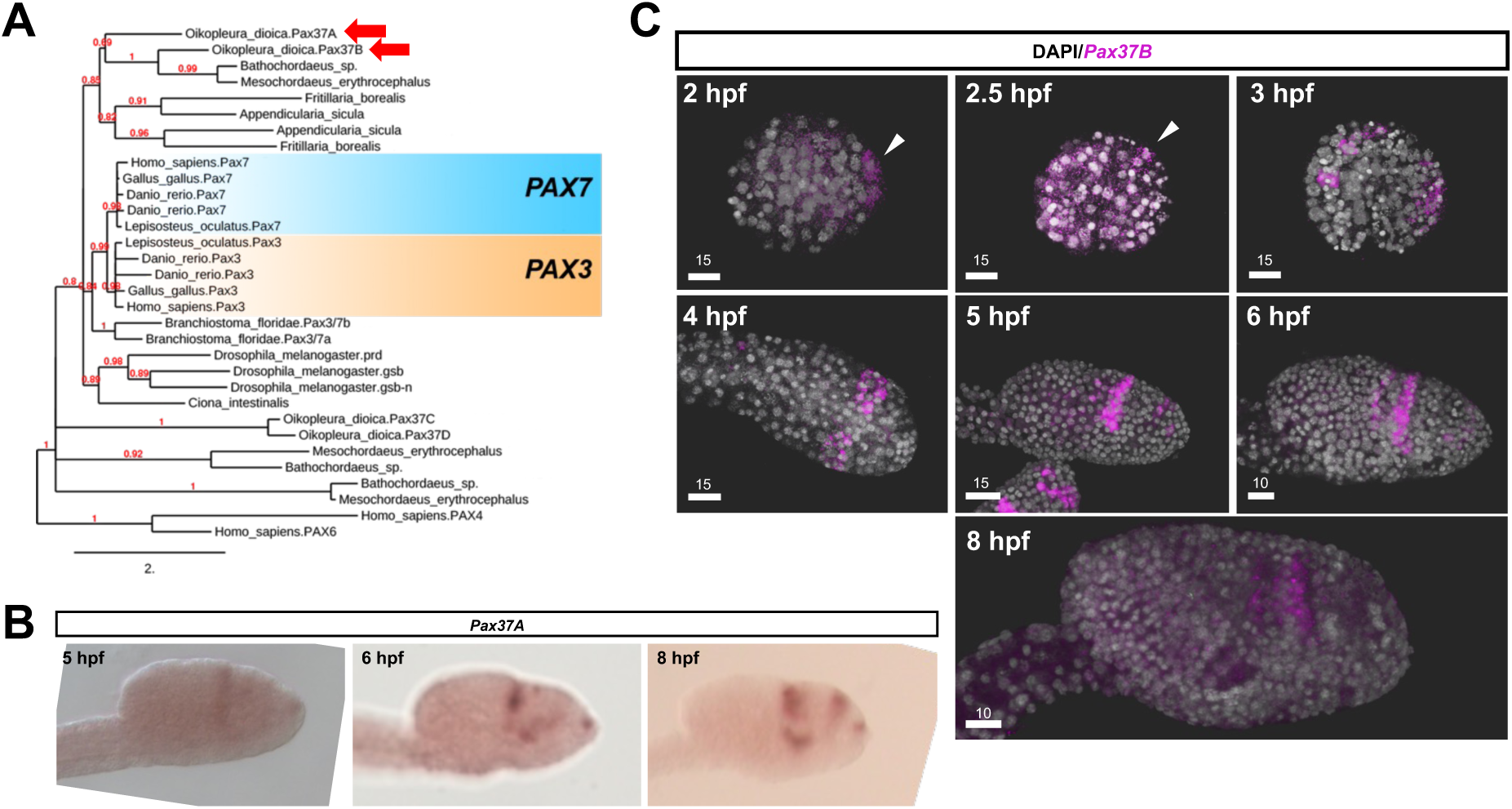
Phylogenetic tree of chordate Pax3/7 genes and expression of *Oikopleura dioica Pax37A* and *Pax37B* at different developmental time-points. A) Phylogeny where *O. dioica* Pax37A and Pax37B genes have been highlighted with red arrows. B) Expression of *Pax37A* at three developmental stages. C) Expression of *Pax37B* during development arrowheads indicate epithelial expression in early stages. Number above scale bar is in μm. Animals are oriented with anterior to the right and dorsal to the top.

### 2.2 *Pax37A* and *Pax37B* have similar but not fully overlapping expression domains

Our focus on Pax37A/B genes is due to their transient expression in the oikoplastic epithelium. With standard ISH (at 5, 6 and 8 hpf) we could observe staining with *Pax37A* probes in the developing posterior Fol, dorsal and lateral developing Nasse/giant Fol cells, anterior Fol and close to the mouth (Fig. 2B). *Pax37B* starts being expressed weakly around 2 hpf on the surface of the embryo (Fig. 2C). At 2.5 hpf staining is still strongest in the epithelium while at 3 hpf (late-tailbud stage) expression in two clusters of nerve cord cells and in cells of the dorsal tail epithelium close to the trunk was observed (Fig. 2C). At this stage the expression in the anterior trunk epithelium has expanded and staining in the developing anterior CNS can be observed (Fig. 2C). Later, at hatching (4 hpf), *Pax37B* expression has started in cells close to where the mouth will form, continued expression in developing anterior CNS, anterior OE developing into the Fol region and still in some nerve cord cells of the tail (Fig. 2C). At 4 hpf no staining could be observed in the tail epithelium. During OE development, between 4 and 6 hpf ^5,6^, expression appear strongest in dividing giant Fol and Nasse cells (Fig. 2C). Later, expression in the giant Fol disappears and it appears in a single row of cells in the posterior Fol (Fig. 2C). Additionally, there is expression in the anterior part of the anterior Fol during this time. Only a few posterior Fol cells show expression until 05:40 (hh:mm) hpf. Subsequently, this row of cells expands (Fig. 2C). At 8 hpf all three rows of Nasse cells and a single posterior Fol row are visible and *Pax37B* positive (Fig. 2C).

To precisely compare the expressions of *Pax37A* and *Pax37B* we used a recently acquired whole larval scRNA-seq dataset (Leon *et al*., *in preparation*). We “re-clustered” epidermal cells expressing either *Pax37A* or *Pax37B* (Figs. 3A and B respectively) and could nicely confirm and refine the ISH and FISH observations (Figs. 2B and C). Both genes are expressed in the anterior Fol developing Nasse and giant Fol cells. Later both genes are expressed in the posterior Fol while expression of *Pax37A* in Nasse cells is lost. During development, *Pax37B* is also expressed in the dorsal part of the anterior ring around the developing mouth. At later stages, as seen in FISH experiments, *Pax37B* is expressed in Nasse cells and posterior Fol cells. Determination of cluster identities was done using known expression pattern of identified marker genes of the respective clusters.

**Figure 3:**
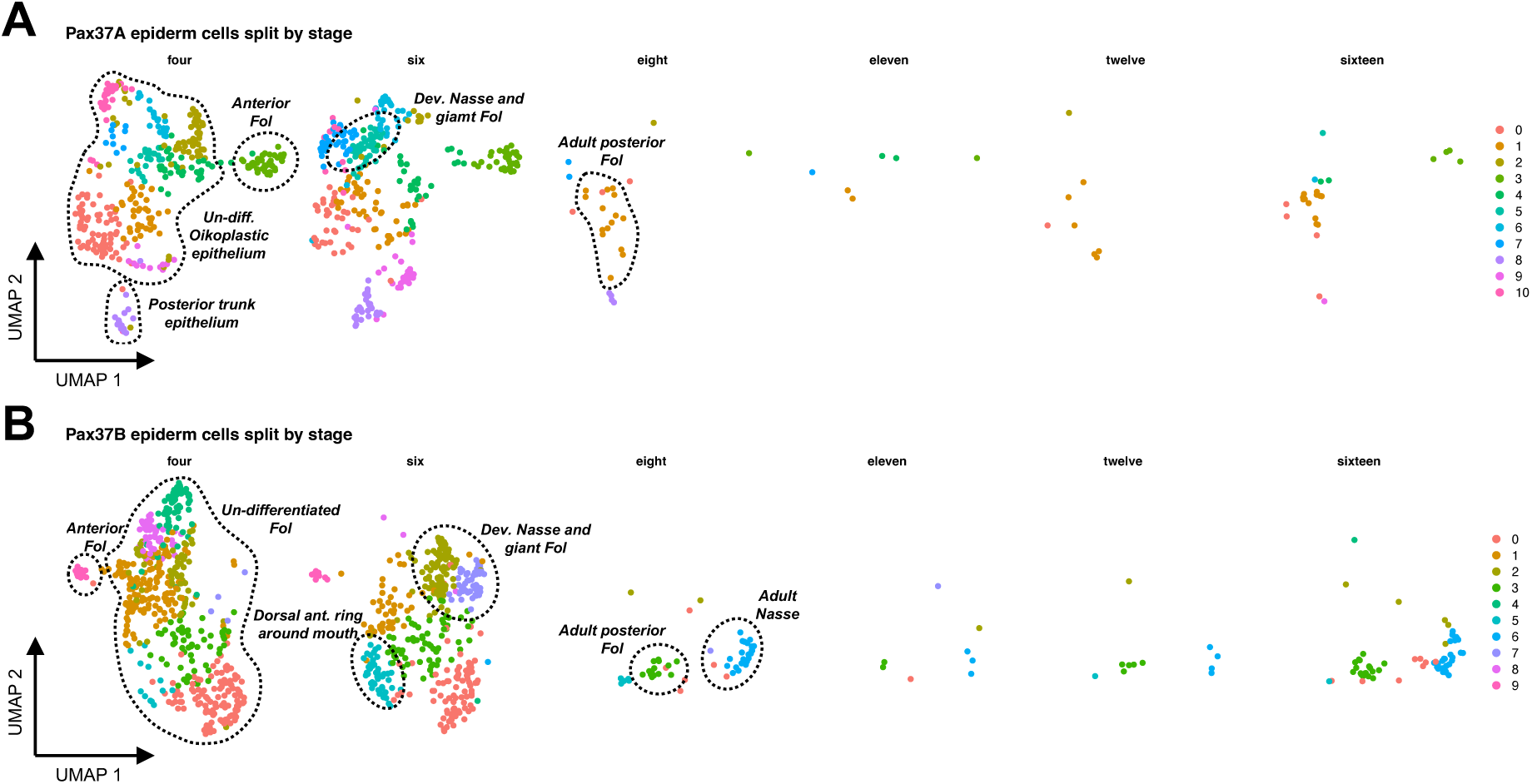
*Pax37A* and *Pax37B* positive cell subsets re-clustered and split by stage of development. A) Re-clustering of *Pax37A+* and B) *Pax37B+* epidermal cells recapitulate the expression patterns observed using ISH and FISH.

### 2.3 Induced mutations show that *Pax37B* but not *Pax37A* is essential for survival

To evaluate the role of Pax37 proteins in the development of the oikoplastic epithelium two stable mutant lines were generated using CRISPR-Cas9: one with a 9 bp change (5 bp insertion and 4 bp substitution) downstream of the paired domain and upstream of the homeobox in *Pax37A* and one with a 4 bp deletion in the paired domain upstream of the homeobox of *Pax37B*. Both mutations resulted in premature stop codons (Fig. 4A).

**Figure 4:**
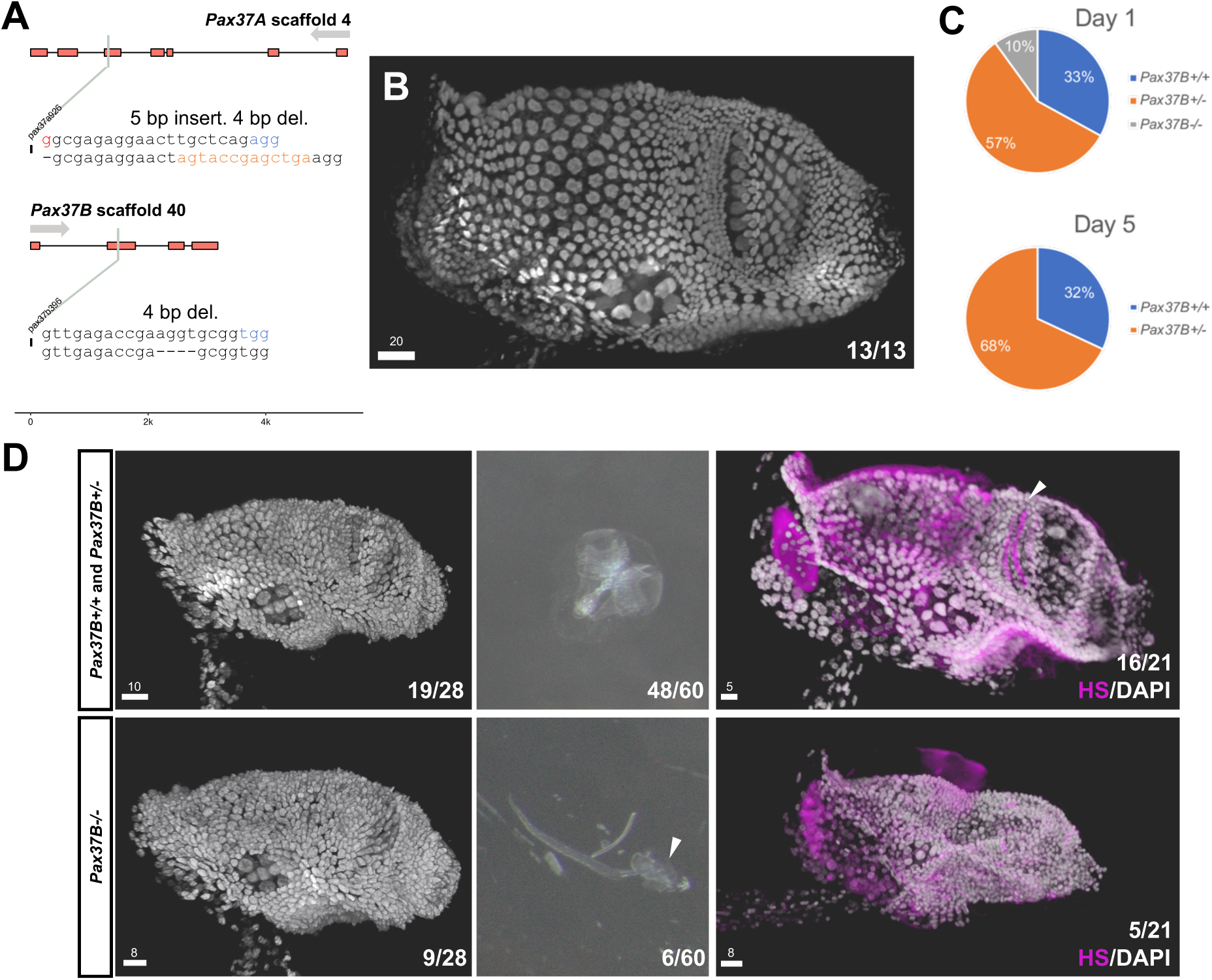
Generated CRISPR mutations and phenotypes in *Pax37A^-/-^* and *Pax37B^-/-^*animals. A) locations of sgRNA targets in the Pax37A and Pax37B gene models and the respective mutations. Upper rows represent the target sequence, and the bottom rows represent the mutation. Blue text represents the PAM sequence and red represent added guanine to the target sequence to aid *in vitro* transcription. B) Representative example of epithelial phenotype in screened animals from crosses of *Pax37A^+/-^*animals. C) Percentage of animals per genotype at different days of collection from crosses of *Pax37B^+/-^* animals. D) Phenotypes of *Pax37B^-/-^* animals compared to their wild-type or heterozygote siblings in the epithelium and house. Fractions correspond to number of imaged animals with that genotype with the phenotype imaged of all genotyped. Number above scale bar is in μm. Animals are oriented with anterior to the right and dorsal to the top.

No obvious phenotype change, either behavioral or in the OE organization, could be observed in the *Pax37A^-/-^* animals when compared to their wild-type and heterozygote siblings at 24 hpf (Fig. 4B). These animals produced fully functioning houses and could be reproduced. In contrast *Pax37B^-/-^*animals died early (at ∼24 hpf only a few are *Pax37B^-/-^* (10 %) and at 5 dpf all of these are gone, see Fig. 4C). Strikingly, at 12 hpf, their OE lacks the characteristic giant Fol cells and is disorganized in the most posterior area of the Fol region. The giant Fol cells and the three rows of highly organized Nasse cells usually seen in animals at this stage appear to be absent (first column in Fig. 4D). Additionally, the anterior Fol area of these animals seems smaller in the *Pax37B^-/-^* animals (first column in Fig. 4D). When mixed animals heterozygous parents were collected at 24 hpf, seven were not inside an inflated house and were unable to inflate house rudiments. Six of them were *Pax37B^-/-^* animals (see second column in Fig. 4D). As a similar house phenotype had been reported when heparan sulfate synthesis was blocked ^2^, we decided to stain offspring from two *Pax37B^+/-^*parents with an anti-HS antibody at 24 hpf. No staining was observed in the food concentrating filter (*fcf*) of six out of 21 animals and five of them were genotyped as *Pax37B^-/-^* animals (third column in Fig. 4D), indicating that sulfation of heparan is reduced in the Fol region and that the *fcf* is not produced in the mutant animals.

### 2.4 *Pax37B^-/-^* animals and failure to differentiate Fol cells

Gene expression studies using mutants began with the identification of differentially expressed genes using whole animal RNA-seq, and the results were then analysed further using our scRNA-seq dataset.

To pinpoint the putative downstream genes affected by knockout of *Pax37B* we performed RNA-seq on embryos at three stages of OE development: 4 hpf, 6 hpf and 8 hpf (Table S1). At 4 hpf we could identify 125 downregulated genes and 103 upregulated genes (padj ≤ 0.05), at 6 hpf we could identify 279 downregulated genes and 453 upregulated genes (padj ≤ 0.05) and at 8 hpf we could identify 457 down-regulated genes and 482 up-regulated genes (padj ≤ 0.05). As *Pax37B* is expressed in several tissues other than the OE, we were not surprised that differentially expressed genes include many predicted to have a function in the central nervous system or in muscles. We observed an enrichment of cell-cycle genes among the downregulated and different pathways relating to metabolism among the upregulated genes at 4 and 6 hpf. At 8 hpf the enriched pathways were different, with the downregulated genes relating mostly to metabolism and the upregulated genes relating to smooth muscle and immune system function (Fig. S1 and Table S2 for gene lists and their best EggNOG hit). *Pax37A* were not among these differentially expressed genes, which fits with our observation of *Pax37A* staining in ISH on larvae at 8 hpf from a cross between *Pax37B^+/**-**^* animals (Fig. S2).

To better discriminate the regulatory cascade that Pax37B is involved in within the OE we queried the above mentioned scRNA-seq dataset of *O. dioica* for stages 4, 6, 8, 11, 12 and 16 hpf. *Pax37B+* epidermal cells were re-clustered using Seurat, and a lineage analysis was performed using Monocle3 (Fig. 5A). This analysis identified 2145 lineage marker genes (q-value < 1e-3, see Table S3). When these lineage markers were compared to the DE gene lists, we observed that the genes that are enriched among the down-regulated lineage markers are mainly involved in processes related to glycosaminoglycans and sulfation while the up-regulated lineage markers mainly are involved in metabolism, cell-cycle, and gene expression (Fig. 5B and 5C). When analyzing the expression of DE oikosins and putative oikosins (identified through Silix blast clustering) among the *Pax37B+* epidermal cell subset we could see that only down-regulated oikosin genes have specific expression in the anterior Fol, differentiated Nasse cells or posterior Fol, while the up-regulated oikosins are not specific to any *Pax37B+* epidermal cells (Fig 5D and 5E).

**Figure 5:**
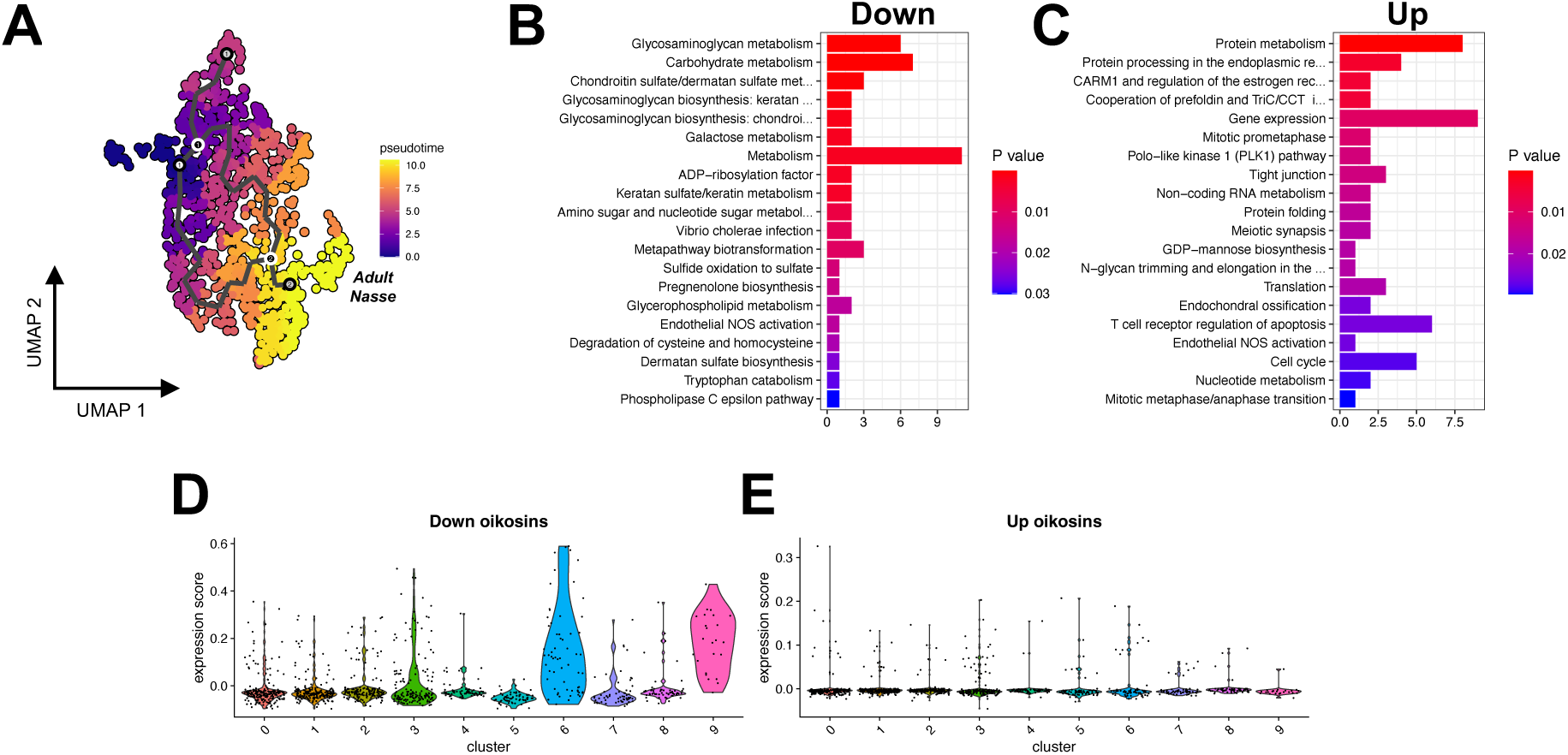
Lineage analysis of the *Pax37B+* epidermal cell subset and expression scores of down- and up-regulated oikosin genes. A) A lineage trajectory of the *Pax37B+* epidermal cells made using monocle3. B) Enrichr pathway analysis of lineage markers that are down-regulated at any of the three stages of bulk RNA-seq show an enrichment of terms related to glycoproteins. C) Enrichr pathway analysis of lineage markers that are up-regulated at any of the three stages of bulk RNA-seq show an enrichment of terms relating to protein metabolism and gene expression but also several cell-cycle terms. D) Expression of down-regulated oikosins and oikosin-like genes identified through silix analysis are mostly restricted to clusters 6 and 9 which represent Nasse cells and anterior Fol cells respectively. E) The up-regulated oikosins and oikosin-like genes identified through silix analysis have very low expression in the *Pax37B+* epidermal cells. Dots in violin plots represent expression score in individual cells of each cluster.

By plotting the expression of the top 20 genes of oikoplastic field specific markers we selected clusters that include cells most likely destined to be Nasse or giant Fol cells (clusters 2, 6, 7 and 8 of the *Pax37B+* epidermal subset) (Fig. S5) and then re-clustered and performed a lineage analysis (Fig 6A, see Table S4). Genes annotated as being transcription factors or having potential transcription factor activity were examined further. When comparing these genes with the DE gene lists, we could find an overlap of nine genes, out of which all but three were upregulated in the mutants. These upregulated transcription factors are mainly expressed early or in the middle of the lineage (6/7) while the downregulated transcription factors are expressed later (2/3) (Fig. 6B). One of the upregulated transcription factor genes had a specific expression in the scRNA-seq dataset and in previous work from our lab ^5^ has been shown to be expressed in the developing Fol, namely *IrxA* (Fig. 6C and D). FISH on larvae from crosses of *Pax37B^+/-^* parents confirmed that the upregulation most likely is an expansion of the expression domain in the Nasse/giant Fol cell precursors (Fig. 6E).

**Figure 6:**
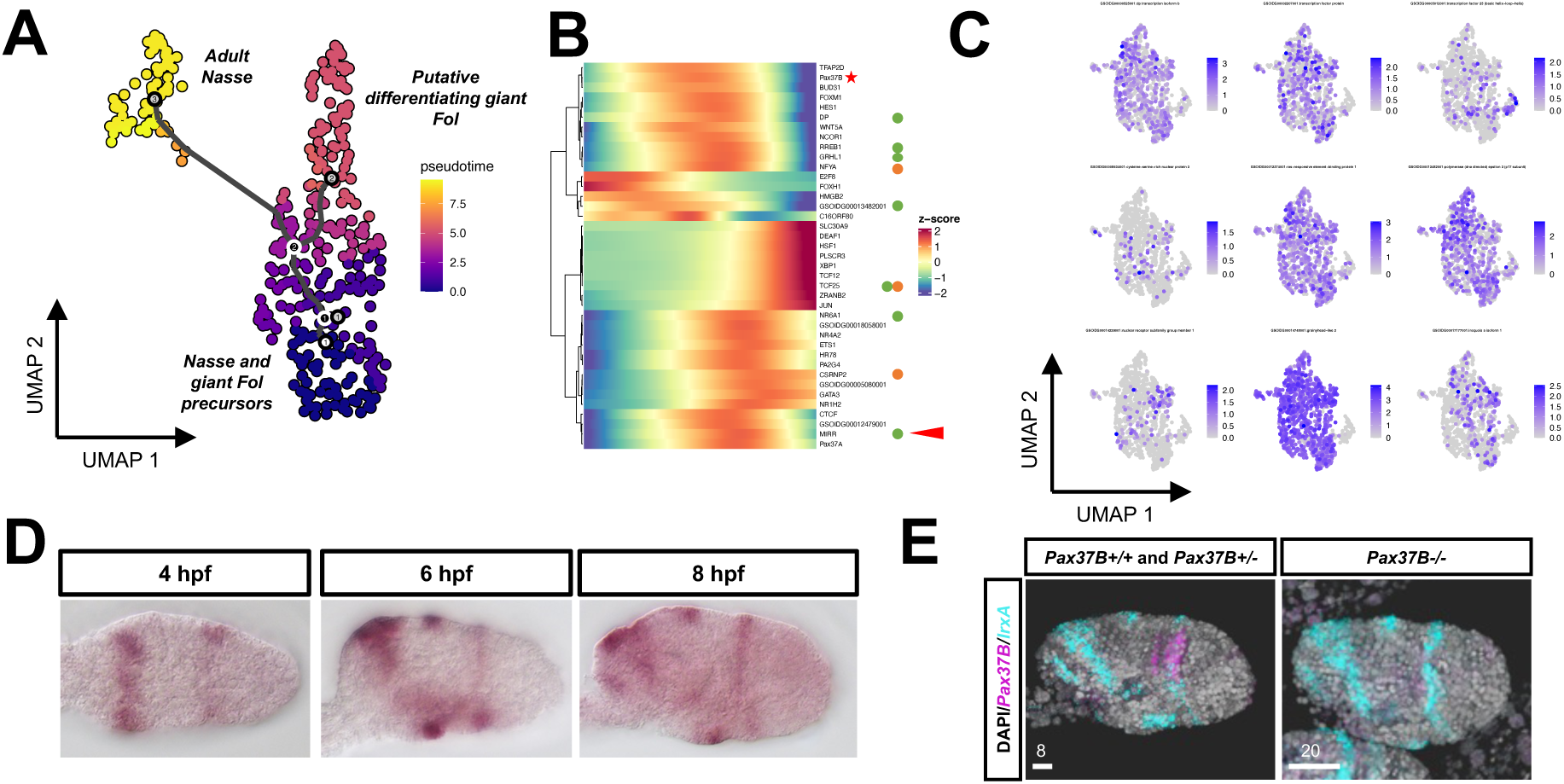
Reclustering and monocle3 analysis of Nasse and putative giant Fol precursor cells. A) monocle3 analysis with trajectory. B) Expression in the trajectory of lineage markers that are related to transcription factor activity. Orange dots represent down-regulated genes and green dots represents upregulated genes. *Tcf25* is up-regulated at 6 hpf and down-regulated at 8 hpf. Red arrow point to *IrxA* which is known for being expressed in the developing Fol. Red star indicate *Pax37B*. C) Expression in the *Pax37B+* epidermal cells of differentially expressed lineage markers that are related to transcription factor activity. D) Expression of *IrxA* at four developmental time-points. E) Expression of *IrxA* in wild-type and heterozygote siblings compared to *Pax37B^-/-^* animals show an expanded expression domain in the developing Fol region. Number above scale bar is in μm. Animals are oriented with anterior to the right and dorsal to the top.

Among the lineage markers of both subsets we find many components of the *O. dioica* Wnt signaling pathway and planar cell polarity pathway ^22^ such as *Wnt5*, *Wnt11d* (annotated as *WNT5A* in our EggNOG analysis but described as *Wnt11d* by Martí-Solans *et al.,* 2021 ^22^), *Ctnnb1*, *Fzd5/8*, *Fzd3/6a*, *Fzd3/6b CK2*, *GSK3*, *Dvl*, *Strabismus/van Gogh*, *RhoA* and *TCF/LEF*. These Wnt signaling pathway components have both similar and different expression among the *Pax37B+* cells (Fig. 7) and, notably, none of these genes are found to be differentially expressed in the mutant animals.

**Figure 7:**
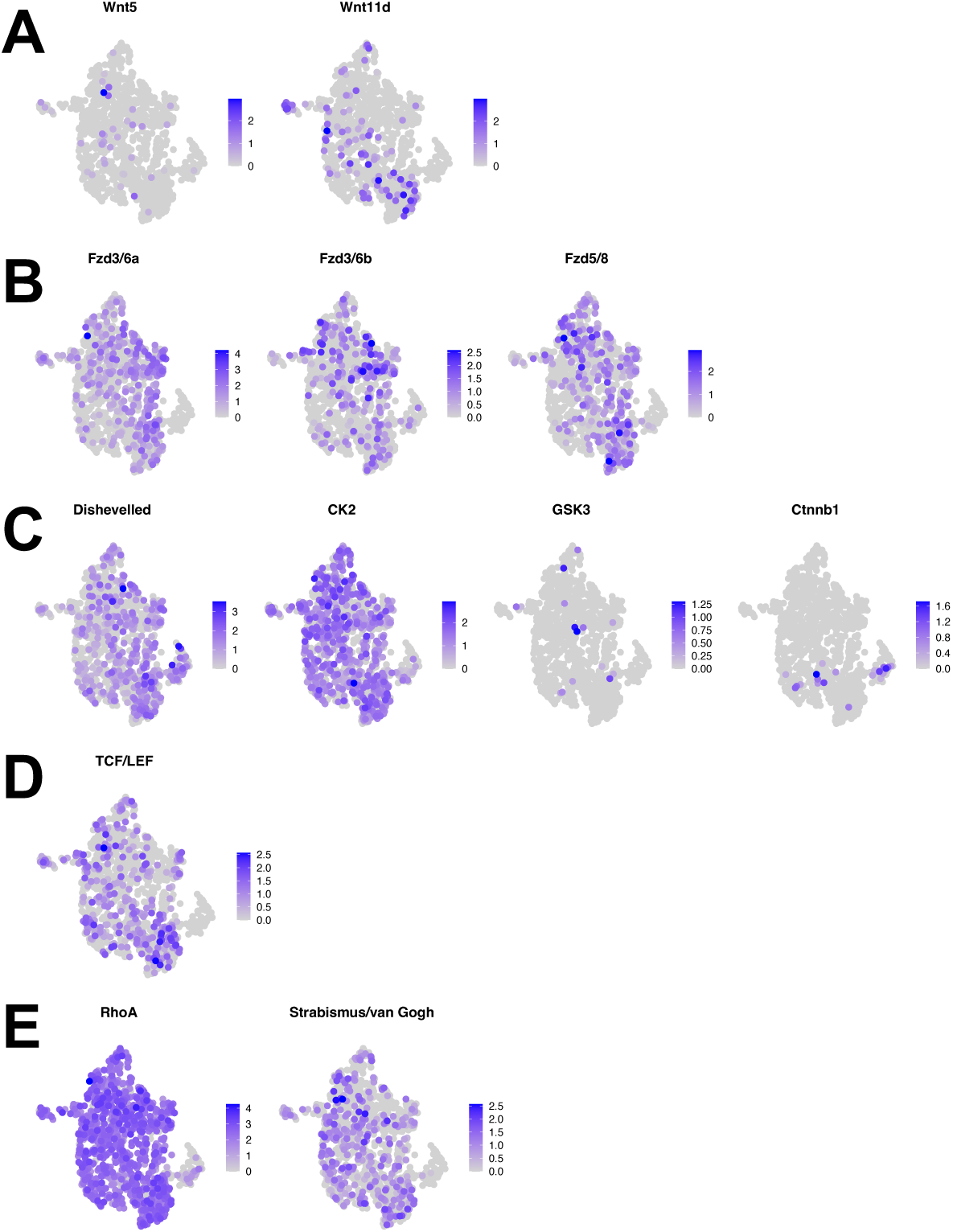
Expression of canonical and non-canonical Wnt signaling pathway components that are lineage makers among *Pax37B+* epidermal cells. A) Expression of Wnt ligands. B) Expression of receptors. C) Expression of intermediate proteins in the signaling cascade. D) Expression of effector transcription factor. E) Expression of planar cell polarity components. All annotations except for *Ctnnb1* were retrieved from ^22^.

### 2.5 Nasse lineage and putative giant Fol specific transcription factors have a similar expression in *Pax37+* epidermal and nervous system cells of *Ciona robusta*

Since *Pax37* have a known expression pattern in the ascidian *C. robusta*, we set out to explore the expression pattern of lineage markers for Nasse and developing giant Fol lineage specific transcription factors. First, we made a dataset of predicted orthologs between *O. dioica* and *C. robusta*. This dataset was then queried to extract the orthologs for Nasse and developing giant Fol cell markers and lineage specific transcription factors. The expression of these orthologs were plotted in *Pax37+* epidermal and a *Pax37+* neuronal cell subsets of the full 10 stages *C. robusta* dataset ^23^. With the data split per stage, we observe a clear division of strongest expression of genes between early and late stages of development, as well as a wide expression in most of the cells (Fig. 8A and B).

**Figure 8:**
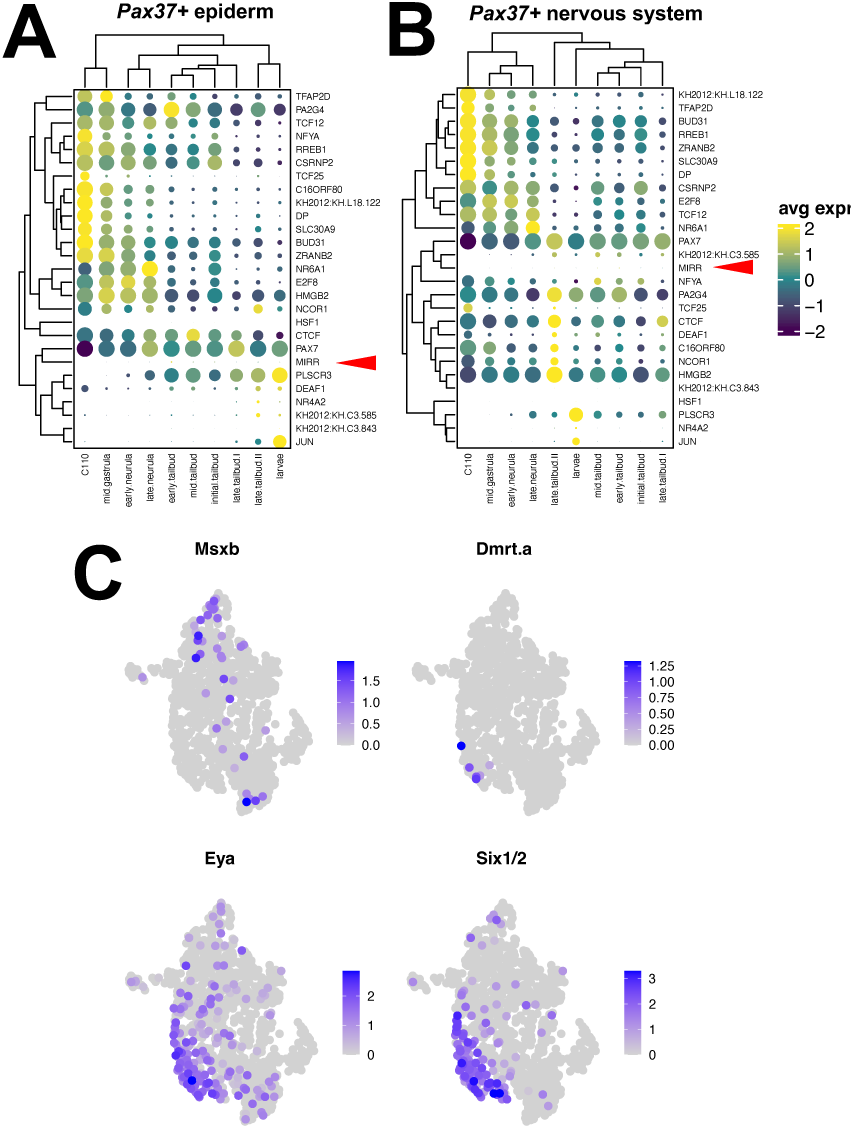
Expression of Nasse and putative giant Fol precursor lineage transcription factors in *C. robusta,* expression of lateral neural plate border marker orthologs in *O. dioica Pax37B+* epidermal cells. A) epidermal and B) neuronal *Pax37+* cells split by developmental stage. Many of the *C. robusta* orthologs have their highest expression already at the initial gastrula stage in both cell types however in the neuronal cells more genes have their highest expression at the late tailbud II stage. A red arrow indicates the expression of the *Ciona* ortholog of *IrxA*. C) Expression of *O. dioica* orthologs for *C. robusta* epidermal neuron markers in the *Pax37B+* epidermal cells.

Interestingly many of the identified transcription factors have their highest expression early during development in both subsets, already at the initial gastrula stage. In the neuronal subset a larger proportion of genes have their highest expression later at the late tailbud II stage. In both datasets the ortholog of *IrxA* has very low to no expression (as indicated by a red arrowhead in Fig. 8A and B). Among the lineage markers for *Pax37B+* epidermal cells we could also identify the ortholog of *C. robusta Msxb* (GSOIDG00001108001:muscle segmentation homeobox) which is a marker for the lateral neural plate border cells that later will develop into the posterior apical trunk epidermal neurons (pATENs) and bipolar tail neurons (BTNs), the ortholog of *Dmrt.a* (GSOIDG00002014001:doublesex- and mab-3-related transcription factor a2) which labels cells that will become aATENs and the palp sensory cells (PSCs) and the orthologs of *Eya* (GSOIDG00002684001:eyes absent 1 isoform 1) and *Six1/2* (GSOIDG00009116001:sine oculis homeobox homolog) which labels cells that will later become aATENs ^18^. When plotted in the *Pax37B+* epidermal dataset a clear division between the expression domains of these genes is seen, where the ortholog of *Msxb* is expressed in the developing Nasse lineage and the orthologs of *Dmrt.a*, *Eya* and *Six1/2* is expressed in the *Pax37B+* cells in the anterior part of the anterior Fol/ring around mouth (Fig. 8C).

When plotting the scaled expression scores of *C. robusta* orthologs of *Pax37B+* epidermal cell cluster markers in the full *C. robusta* 10 stages dataset we observe higher expression scores in epidermis, nervous system and muscles and mesenchyme (Fig. 9A). When plotting the expression scores of the orthologs of DEGs of different stages in the full dataset we see a similar pattern (Fig. 9B). In the larval CNS dataset, we observe that most *Pax37B+* epidermal cluster markers have the highest expression in the anterior CNS in cells such as PSCs, collocytes, aATENs, CESNs, neurohypophysis primordium and pATENs but also in glial cells compared to the orthologs of *O. dioica* neuronal cluster markers (Fig. 9C). This observation is also true for markers of other Fol regions and the giant Eisen region (Fig. S6). When investigating the expression scores of the DEGs in the larval CNS dataset we observe a similar pattern (Fig. 9D).

**Figure 9:**
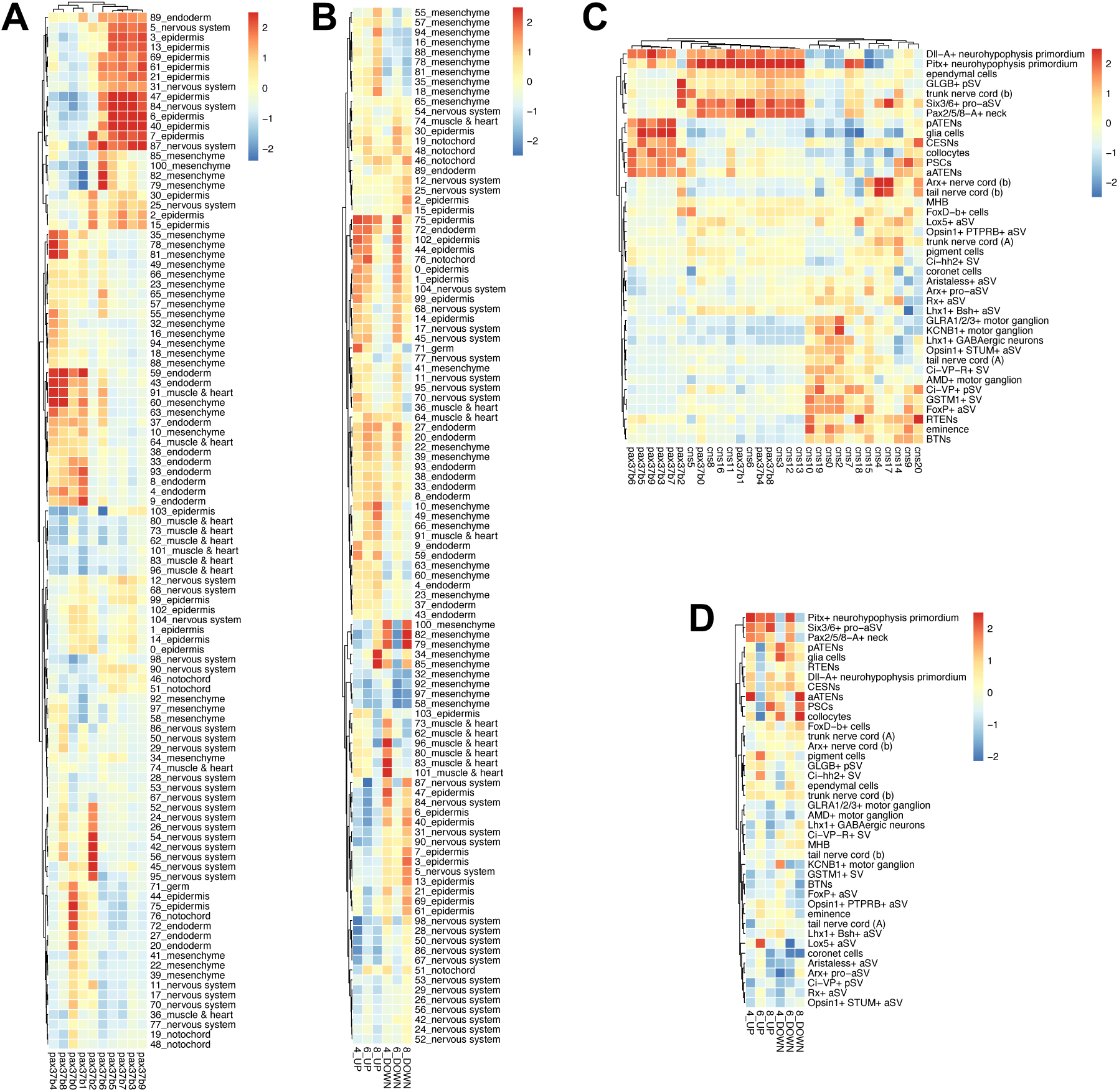
Expression scores of *Pax37B+* epidermal cell cluster markers as well as up and down regulated genes at 4, 6 and 8 hpf. A) The expression scores of Pax37B epidermal cell cluster markers are highest in the full *C. robusta* dataset epiderm and nervous system clusters. B) The expression scores of down regulated genes at 4, 6 and 8 hpf have the highest expression in epiderm, nervous system, muscle & heart and mesenchyme clusters. C) The expression scores of *Pax37B+* epidermal cell cluster markers in the larval CNS dataset of *C. robusta* are highest in the anterior nervous system, especially in PSCs, collocytes, aATENs, pATENs and CESNs, as well as glial cells. In this plot *O. dioica* putative CNS clusters from the full dataset also were included. D) The expression scores of down regulated genes at 4, 6 and 8 hpf have in the larval CNS dataset the highest expression in the anterior nervous system, collocytes, aATENs, pATENs, as well as glial cells.

To investigate the neuronal and epidermal properties further we plotted the *C. robusta* orthologs of Nasse cell markers (cluster 6 in the epidermal *Pax37B+* reclustering) in the full and in the larval CNS datasets. This revealed a subset of genes that contribute the most to the observed similarity. These genes have their highest expression in epidermal and nervous system clusters in the full dataset and in certain nervous system cells in the larval CNS dataset (Fig. 10A and B). When investigating the annotations of these *C. robusta* genes, we find that they are involved in extracellular matrix, cytoskeletal components, and focal adhesion, not necessarily specific for epithelial or neuronal function (Table S5). Similarly, when investigating all *Pax37B+* epidermal clusters we observe no, or few genes involved in mature neuron signaling and function, however one homolog of muscle specific acetylcholine receptors, *CHRNG*, is expressed in adult Nasse cells.

**Figure 10:**
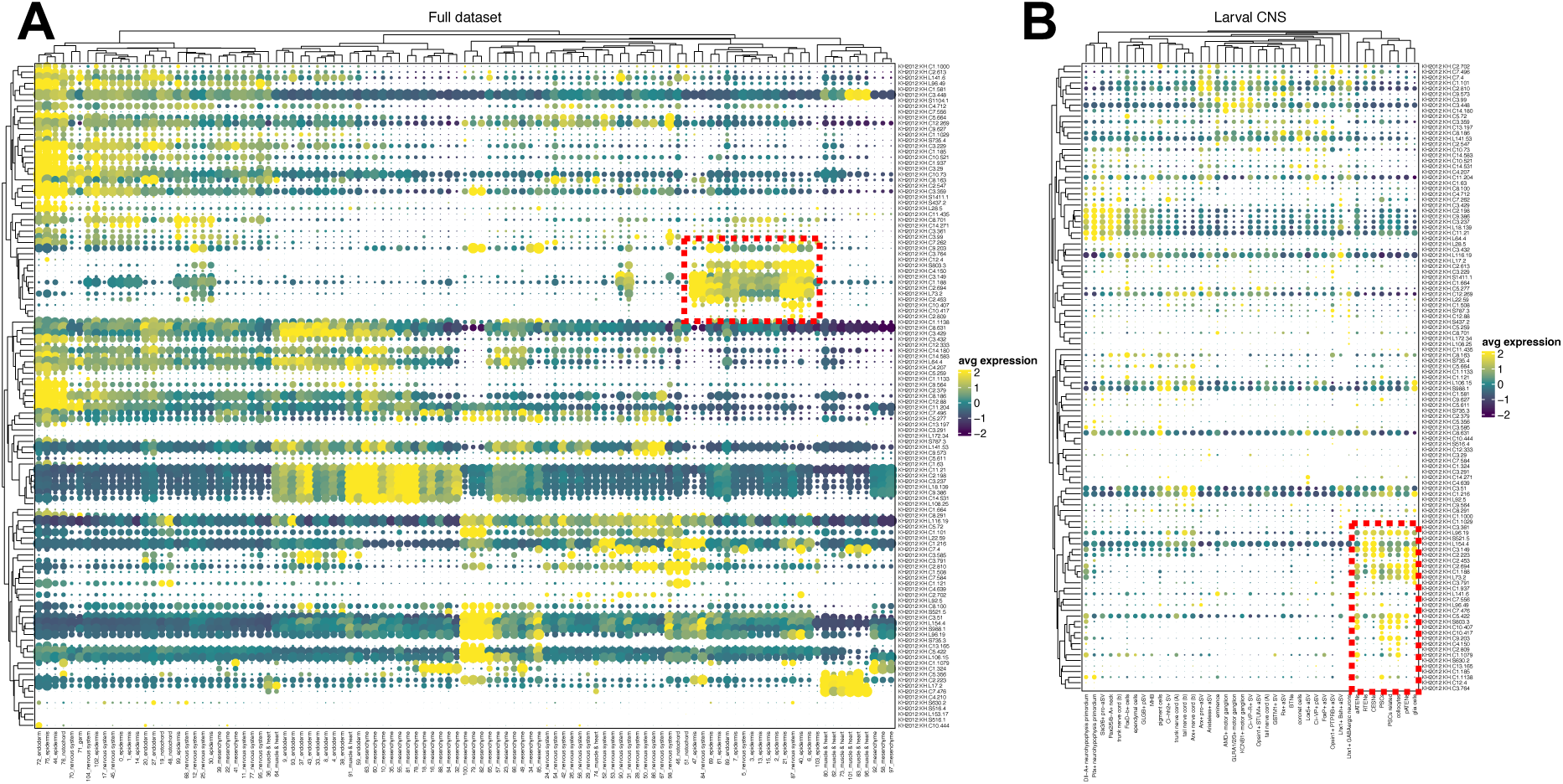
Expression of *Ciona robusta* orthologs of adult Nasse cell markers in the full *Ciona robusta* dataset (A) and the larval CNS dataset (B). Red boxes highlight genes that are specific to epiderm and/or nervous system clusters in (A) and epidermal and glial cell clusters in (B).

## 3. Discussion

### 3.1 *Pax37B* is essential for the development of Fol’s oikoplast

In this work we have shown the important role of Pax37B in the development of Fol’s oikoplast. This contrasts with Pax37A which do not give any observable phenotype when knocked out. Despite the co-expression of *Pax37A* and *Pax37B* in some cell types of Fol’s oikoplast Pax37A does not compensate the loss of Pax37B and *Pax37B* mutants eventually die at ∼24 hpf. An important role for Pax37 has been reported for the proliferation and differentiation of tissues originating from the lateral neural plate (ascidian tunicates) or neural plate border and later neural crest (vertebrates) but also for regulating neural tube closure (both tunicates and vertebrates) ^12,13,15,24^. In *O. dioica*, the neural plate or the closure of the neural tube have not been observed ^19^. However, due to the conserved role of Pax37 in this specific process in ascidians and vertebrates, Pax37A and/or Pax37B most likely have a similar role in *O. dioica*.

During late embryonic and early larval development *Pax37A* and *Pax37B* are both expressed in specific domains of the dorsal trunk OE (Mikhaleva *et al.,* ^5^; Fig. 2B and C). Their expression domains are partially overlapping, especially in the posterior Fol region (Fig. 2B and C). The lack of a phenotype in the OE of *Pax37A^-/-^*animals may be the result of a compensation by Pax37B in these cells. On the other hand, when *Pax37B* is knocked out we can observe a severe phenotype in the OE in regions where we observe no or little overlap with *Pax37A* expression such as the giant Fol and Nasse cells. This indicates that *Pax37A* is not able to compensate for the loss of *Pax37B* in these cells.

### 3.2 *Pax37B* is responsible for controlling proliferation and differentiation of cells in Fol’s oikoplast

Due to the broad and conserved roles of Pax37 genes in chordates, where they are involved in the proliferation and differentiation of precursor cells, we set out to investigate how novel the mode of epidermal expression of *Pax37B* and *Pax37A* is in *O. dioica*.

Based on our observations in this study we hypothesise that Pax37B plays a role in the proliferation and differentiation of oikoplastic epithelium cells for three main reasons: 1) In the *Pax37B^-/-^* animals, genes that relate to cell-cycle pathways are enriched among the genes that are down-regulated at the earliest stages. At later stages, oikosins and proteins with similarity to known oikosins expressed in *Pax37B+* OE regions are down-regulated; 2) The down-regulation of lineage markers of *Pax37B+* epidermal cells that are genes involved in sulfation and our observation of reduced staining with anti-heparan/heparin sulfate antibody in the *fcf* and failure to inflate the house rudiments like what has been observed in previous studies when blocking this pathway with inhibitors ^2^; 3) Among the *Pax37B^-/-^* animals we observed an increase of expression of two giant Fol specific oikosins, *oikosin1* and *oikosin14* in the bulk RNA-seq data. In FISH experiments these genes are, in the *Pax37B^-/-^* animals, expressed in what appears to be one cell on one side of the animal (Fig. S7).

Additionally, the shared transcription factors between the up-regulated genes and the lineage markers are mostly expressed early in the lineage while the shared downregulated transcription factors are either expressed late after differentiation or involved in cell type specification. Most of the differentially expressed transcription factors have a broad expression in almost all *Pax37B* positive cells while one, *IrxA*, is upregulated and expressed in a distinct subset of cells both in the scRNA-seq data, ISH and FISH. *IrxA* is expressed in developing Nasse and giant Fol precursors (Fig 5D; Mikhaleva *et al.,* ^5^) and the upregulation of *IrxA* observed in the RNA-seq data was confirmed using FISH in *Pax37B^-/-^* animals (Fig. 6E). These observations lead us to propose that Pax37B in combination with IrxA is required for the differentiation and development of Nasse and giant Fol cells. We were thus far unable to obtain a mutant line for *IrxA* and therefore ignore the phenotypic effect of its loss.

### 3.3 Wnt signaling is a candidate activator of *Pax37* expression in the developing Fol’s oikoplast

By investigating the *Pax37B+* epidermal cell and Nasse and putative giant Fol precursor lineage markers we identified multiple components of the canonical and non-canonical Wnt signaling pathway that were previously been annotated in *O. dioica* ^22^. None of these components are differentially expressed in our mutant animals. The canonical Wnt pathway is involved in specification of the neural plate border of vertebrates where it activates *PAX3* expression ^16^. The lack of change in expression of these components in the mutant animals and the expression pattern of *Wnt11d* in the same area as *Pax37B* expression at 3 hpf (Fig. 2B and Martí-Solans *et al.,* ^22^) lead us to propose that Wnt signaling may activate *Pax37* expression in the developing Fol’s oikoplast of *O. dioica*.

### 3.4 Cells of the filter-producing fields of the OE have a neuroepithelium-like signature

The morphology of the trunk epidermis in *C. robusta* and *O. dioica* differ greatly. While the *C. robusta* epidermis is rather uniform with some epidermal sensory neurons the *O. dioica* trunk epidermis has numerous specialized fields of secretory cells producing different parts of the house. Since *Pax37* is expressed in the dorsal epidermis of ascidian tunicates as well as in tissues that will develop into the different apical trunk epidermal neurons ^13,14,18^ we thus hypothesized that the Fol cells might derive from such tissues. This led us to investigate the possibility that the cells that express *Pax37A/B* in *O. dioica* share ancestry with Pax37 positive epidermal/neuronal cells of *C. robusta*.

*Pax37* is already expressed at the gastrula stage of *C. robusta* in cells that will develop into several different tissues such as muscle, epidermis, and nervous system. During neurulation, *Pax37* is mainly expressed in the lateral plate ectoderm ^13^. In *C. robusta*, precursor cells for some of the epidermal neurons express Wnt signaling components ^17^ and *Pax37* ^18^. In our dataset we find in addition to Wnt pathway genes, orthologs of *C. robusta* apical trunk epidermal neuron fate markers, *Dmrt.a*, *Msxb*, *Six1/2* and *Eya*, which are also involved in neuronal fate determination. These are also among the lineage markers and are expressed in different subsets of the *Pax37+* epidermal cells from *O. dioica*. Nasse and putative giant Fol precursor cells express the ortholog of *Msxb* but none of the others while the other genes are expressed in cells anterior of the anterior Fol. This would suggest that distinct cell types of the Fol region share a gene regulatory network (GRN) that governs the development of some of the larval neurons in *C. robusta*. To investigate this further we plotted the expression scores of *C. robusta* orthologs of *Pax37B+* epidermal cell cluster markers and DEG genes in the full and larval CNS *C. robusta* scRNA-seq datasets. This showed that these markers and the DEGs have a higher combined expression score in both epidermis and a few subsets of neurons further indicating that these cells have a neuroepithelial-like signature. However, these shared genes are not markers of mature neurons suggesting specialization in *O. dioica*. Interestingly adult Nasse cells express a nicotinergic acetylcholine receptor possibly indicating that Nasse cells are receiving cholinergic input to control their secretion, however this must be investigated further through functional experiments.

### 3.5 Conclusions

The work presented here shows that the Pax37 proteins indeed play a major role in the genesis of the house, a major innovation that evolved in larvaceans. They control the proliferation and differentiation of cells that becomes the anterior anterior Fol, giant Fol, Nasse and posterior Fol in the oikoplastic epithelium. In these cells, Pax37B plays the main role, while the role of Pax37A is unclear and may be compensated by an active Pax37B in the *Pax37A* mutant. Transcriptional profiles indicate the presence of GRN components recognized in certain *C. robusta* neurons in *Pax37B+* epidermal cells, their lineage markers and adult Field specific markers. Taken altogether the present findings provides support for a “hybrid cell identity” of the Fol cells of *O. dioica*, being epidermal cells under control of a GRN operating in some ascidian neuronal subtypes. Interestingly, one ascidian neuron type that shares high similarity with the *O. dioica* Nasse cells is that of collocytes, cells which are secreting the glue used for attachment to the substrate before the metamorphosis of ascidians. Finally, these findings provide support to the hypothesis that co-option of existing GRNs and components of GRNs play an important role in evolution of novel features ^25^.

## 4. Materials and Methods

### 4.1 Animals

The *O. dioica* used in this study, collected in the coastal waters of Bergen, Norway, had been cultured for several generations at 13°C in the Michael Sars Centre Larvcean facility before experiments. All microinjections were performed in a temperature-controlled room at ∼18°C and using filtered artificial seawater (FASW).

### 4.2 identification and phylogenetic analysis of Pax3/7 genes

Known amino acid sequences of Pax3/7 genes were collected from NCBI, Uniprot and Ensembl. *O. dioica* sequences were then used in blast searches against additional larvacean genome assemblies ^26^ and annotated manually as well as using genBLAST ^20,21^. Sequences were aligned using the http://www.phylogeny.fr web server with MUSCLE (using standard settings) and a phylogenetic tree was made using PhyML (with aLRT, SH-like, branch support option).

### 4.3 RNA extraction, RACE reactions, target identification, sgRNA synthesis and injections

RNA was extracted from ∼50-100 animals of different stages were collected in a small amount of FASW and snap-frozen in liquid N_2_ and stored in -80°C until extraction. Total RNA was extracted using the Nucleospin® RNA XS kit (Macherey-Nagel, Cat no: 740902.50S), with the on-column rDNAse step, and eluted in 10 uL RNAse free H_2_O. Concentration of extracted RNA was measured on a Nanodrop and integrity was checked on a 1% agarose gel stained with Sybr Gold gel stain. RNA extracted were used in 5’ RACE ready cDNA reactions using the SMARTer® 5’/3’ RACE Kit (Clonetech, Cat. Nos. 634858, 634859). Primers are provided in Table S7.

The sequences from the RACE reactions were used to identify suitable targets for sgRNAs. Putative target sites for sgRNAs located in the region 5’ of or within the homeobox region of both genes were selected using a custom python script identifying microhomologies of 4, 5 or 6 bp located at most 8 bp apart ^5,6^. Among these those that had a suitable PAM sequence were investigated further to avoid SNPs in the target sequence. Three sgRNAs were designed and tested (See Table S7 for oligos used). The identified sgRNAs were synthesized and injected into unfertilized oocytes. The injection mix contained: 400ng/uL sgRNA, 800ng/uL EnGen® Cas9 NLS (NEB, Cat. No: M0646T). A volume of ∼1/4-1/5^th^ of the egg diameter was injected into the eggs and they were then left to incubate for 2 hours before fertilization. For each batch of eggs, several eggs were left un-injected on the injection stage and collected as un-injected controls that were fertilized at the same time as their injected siblings. Embryos were then collected (embryos injected with the same sgRNA were pooled into one tube and their un-injected control eggs pooled into a separate tube) and snap-frozen in Liquid N_2_ at 4 hpf for gDNA extraction and genotyping using High Resolution Melt analysis (HRM) and later confirmed using PCR followed by cloning and sanger sequencing or amplicon sequencing.

Initial Sanger sequencing experiments resulted in the following efficiencies of the best sgRNAs: pax37b396 could edit the desired locus in 15.4% of the clones (6/39) and pax37a926 could edit the desired locus in 7.1% of the clones (4/56). These were the sgRNAs selected for further experiments.

### 4.4 Generation of stable mutant lines

The most efficient sgRNAs were used for injection into new batches of eggs ^27^ that were after successful fertilization collected for culture following the standard protocol ^28^. At day 6 mature animals were screened using qPCR with HRM analysis. 20 uL of sperm (spawned in 1 mL FASW) or ∼20 2 hpf embryos fertilized with wildtype sperm in 20 uL FASW were collected in individual PCR tubes. Samples were centrifuged for 10 min, 10500 rpm at 4°C and supernatant were removed. 5 uL proteinase K mix from the ARCTURUS® PicoPure® DNA extraction kit (Applied Biosystems, Cat no. KIT0103) was added to each sample and they were incubated for 30 min at 65°C followed by 5 min at 95°C. 1 uL of the extracted genomic DNA were used for qPCR and HRM analysis using the IQ SYBR® Green Supermix kit (BioRad, Cat no. 1708882). Mutation positive male sperm were used to fertilize wildtype eggs which were saved in culture beakers and raised until next spawn. At the next spawn the procedure was repeated to identify heterozygote animals that in turn were crossed with wildtype partners, however at this time the bodies of spawned females were screened instead of fertilized eggs. The resulting lines of heterozygotes were, after validation of suitable mutations by sequencing, used in the subsequent experiments.

### 4.5 Imaging of OE and genotyping of animals

Animals from single crosses between heterozygote parents were raised until 6, 9 and 12 hpf, at 12 hpf the different fields of the oikoplastic epithelium have been shown to be fully established ^5,6^, and collected and fixed overnight at 4°C in 4 % paraformaldehyde. Fixed animals were washed 2 x 30 min PBSTEG, 2 x 5 min PBSTE, blocked for 1 hour at room temperature in Blocking solution (3% BSA; 0,02% sodium azide in PBSTE) under agitation, washed 1 x 30 min at room temperature in PBSTE-1% BSA supplemented with 2% phalloidin-AF488 (Invitrogen A12379), washed 4 x 30 min at room temperature in PBSTE-1% BSA and washed 2×30’ at room temperature in PBSTE. Post washes the animals were stained for nuclei with hoechst (1 ng/uL in PBS) for 10 min or DAPI (1/4000 in PBS) for 2h at 4°C followed by 3 x 5 min washes in PBS + 0.02% Sodium Azide. Before imaging the animals were mounted individually in 1.5 % low melt agarose in PBS. All confocal stacks were acquired using Leica SP5 or Olympus FV3000 point scanning confocal microscopes. 3D reconstructions were made using Imaris Viewer 9.7.2 (https://imaris.oxinst.com/imaris-viewer). After imaging animals were collected in individual PCR tubes and genotyped as follows: 10 uL proteinase K mix from the ARCTURUS® PicoPure® DNA extraction kit was added, incubated at 65°C for 3 hours then 95°C for 5 min. 2 uL template were used for PCR followed by sanger sequencing. The *Pax37A* animals from heterozygote parents were imaged mounted on a glass slide and thus not genotyped.

### 4.6 Whole mount *in situ* hybridization and fluorescent *in situ* hybridization

Probes for whole mount ISH and FISH were prepared with RT-PCR using primers listed in Table S7 followed by ligation into pGEM-T Easy vectors and transcribed using a DIG RNA labeling kit (Roche, Cat no. 11175025910), some probes were made using flourescein labeling mix (Roche, Cat. No 11685619910). Plasmids for *Pax37A* and *Pax37B* probes was kindly provided by Dr. Yana Mikhaleva. ISH and FISH were performed as described in Mikhaleva *et al.,* 2018 ^5^ with the difference that the TSA-Cy5 and TSA-Fluorescein staining steps was done at 4°C. Additionally, probe concentrations and hybridization temperatures were adapted to each probe, but were usually around 1:400 dilutions in hybridization buffer and incubated at 65°C. DAPI staining was done by incubating the samples in DAPI diluted 1:4000 in PBS at 4°C for two hours. Slides were mounted in 80% glycerol in PBS. FISH samples were imaged as described in section 4.4.

Negative Pax37B staining was used as a marker for *Pax37B^-/-^* animals in double FISH experiments.

### 4.7 Whole mount immunohistochemistry

Whole mount IHC for heparan/heparin sulfate (MAB204, Millipore) was performed in 24 hpf animals as described in Hosp et al., 2012 ^2^. After IHC animals were stained with DAPI as described above. Samples were imaged as described in section 4.5.

### 4.8 RNA-seq samples and preparation and analysis

Animals from crosses between *Pax37B^+/-^* parents were collected individually in 1 uL FASW in PCR tubes and snap frozen in liquid nitrogen at 4 hpf (hatching), 6 hpf and 8 hpf. 10 uL of extraction buffer (RA1) mixed with TECP (from the Nucleospin® RNA XS kit) was added to each sample and they were spun down at 10500 rpm for 2 min then vortexed and spun down quickly to collect all at the bottom of the tube. 1 uL of the lysed samples were then used in genomic DNA extraction for genotyping. The rest was snap frozen in liquid nitrogen for later RNA extraction. Genomic DNA was extracted by adding 5 uL proteinase K mix from the Arcturus® PicoPure® DNA extraction kit and then incubating the samples for 3 hours at 65°C and then 95°C for 10 minutes. 1 uL of the genomic DNA was used for PCR with the Advantage® 2 PCR kit in 50 uL reaction volume. The resulting PCR products subjected to sanger sequencing. Identified wildtype and homozygote mutant samples were pooled in pools of three animals each, resulting in two wildtype pools and two mutant pools at 4 hpf, two wild type and three mutant pools at 6 and 8 hpf. RNA from these pooled samples were extracted following the Nucleospin® RNA XS kit protocol, with the on-column rDNAse step, and eluted in 10 uL RNAse free H_2_O. The quality of the extracted RNA was investigated using an Agilent bioanalyzer. The extracted RNA was used for cDNA synthesis, amplification and sequencing library generation following the New England Biolabs (NEB) protocol for low input RNA (NEB, Cat no #E6420). The quality and quantity of the resulting library was analyzed on an Agilent bioanalyzer. Initial sequencing was performed using an illumina MiSeq machine 2×300 bp reads generating on average 4.7, 2.5 and 2.6 million reads per library per end for 4, 6 and 8 hpf libraries respectively. To achieve higher sequencing coverage, we re-sequenced the 6 and 8 hpf libraries in a single run of an illumina NextSeq 500 machine at the Norwegian Sequencing Centre in Oslo (2×150 bp mid-ouput mode) generating on average 18.6 million reads and 17.3 million reads per library and end for the 6 and 8 hpf samples respectively.

Sequencing reads were checked for quality using FastQC 0.11.5 and trimmed for sequencing adaptors using the recommended protocol from NEB using flexbar 3.4.0 ^29,30^. No quality trimming was performed. Reads were aligned against the annotated transcript sequences from the *Oikopleura dioica* genome assembly version 3.0 using kallisto 0.46.0 ^31^. The output from kallisto were analyzed using DESeq2 in the DEBrowser ^32^ package in R. Data were filtered for >5 mapped reads in at least two of the four/five analyzed pools and that show an adjusted p-value of ≤0.05 and a fold change of greater than two.

All peptide sequences in the file Oikopleura_peptide_v1.0.fa (retrieved from http://www.genoscope.cns.fr/externe/Download/Projets/Projet_HG/data/annotation/) was annotated using the EggNOG-mapper v1 server (http://eggnogdb.embl.de/#/app/emapper).

### 4.9 scRNA-seq data analysis of epidermal subsets

Using Seurat 4.1.2 ^33^, a subset for epidermal cells was selected from single cell RNA-seq data set of dissociated whole animals 4, 6, 8, 11, 12 and 16 hpf produced in our lab (Leon *et al.*, in preparation) by the expression of epidermal marker genes such as epidermal homeobox transcription factors and oikosins ^2,5^. This subset was re-clustered and markers for the clusters representing giant Fol, anterior Fol and giant Eisen were also collected.

The *Pax37B* and *Pax37A* subsets were created by selecting cells with more than zero expression of the respective genes from the epidermal subset. These were re-clustered and analyzed further. A lineage analysis was performed for the *Pax37B+* epidermal subset using Monocle3 ^34–37^ and the 4 hpf libraries as a root. Lineage markers with a q-value <1e-3 was then exported for further analysis. An additional subset, re-clustering and lineage analysis was done for clusters that have elevated expression scores for the top 20 adult Nasse and giant Fol specific markers (Fig. S4).

### 4.10 Functional enrichment analysis

Best match gene name reported by EggNOG for each gene in each gene set were used in Enrichr^38–40^ runs against the BioPlanet 2019 pathways. Enriched pathways were plotted in R.

### 4.11 Comparisons between Oikopleura dioica and Ciona robusta

By using Orthofinder ^41^ we identified the best ortholog match between genes in *Oikopleura dioica* and *Ciona robusta* (*intestinalis type A*) (KH2012 assembly). This dataset was then used to find *C. robusta* orthologs for *O. dioica* marker genes. The expression matrices and annotations for the *C. robusta* scRNA-seq dataset was downloaded and the expression scores of orthologs of *O. dioica* marker genes were plotted using Seurat 4.1.2.

### 4.12 Data availability

The data presented in this manuscript has not been deposited in any public repository but is made available upon request to the corresponding author. All scripts used to create the datasets and plots are provided as a supplementary R notebook.

## Supporting information

R notebook with scripts used in this manuscript

Alignment used for the phylogenetic tree

## 5. Acknowledgements

Thanks to Anne Aasjord and Kjerstin Nilsen Nøkling for help with animal care. Thanks to Yana Mikhaleva for valuable input regarding the epithelium of *O. dioica.* This project was supported by a grant for the Sars Centre core budget (NFR grant 234817 “Sars International Centre for Marine Molecular Biology Research, 2013-2022”) and a grant to D.C. (NFR grant 250005 “accelerated evolution in chordates and the origin of larvaceans”), both from the Research Council of Norway.

## 6. Supplementary files

Supplementary_tables.xlsx – compilation of all supplementary tables S1-S7.

Supplementary_figures.pdf – compilation of all supplementary figures S1-S6.

Pax37_alignment.txt – the alignment used to make the phylogenetic tree in Fig. 2A.

Notebook_for_generating_analyses_and_plots_for_manuscript.nb.html – R notebook containing all scripts used to generate the datasets and plots presented in this manuscript.

