## Supplementary figures S1-S6 for "Pax37 gene function in *Oikopleura dioica* supports a neuroepithelial-like origin for its house-making Fol territory"

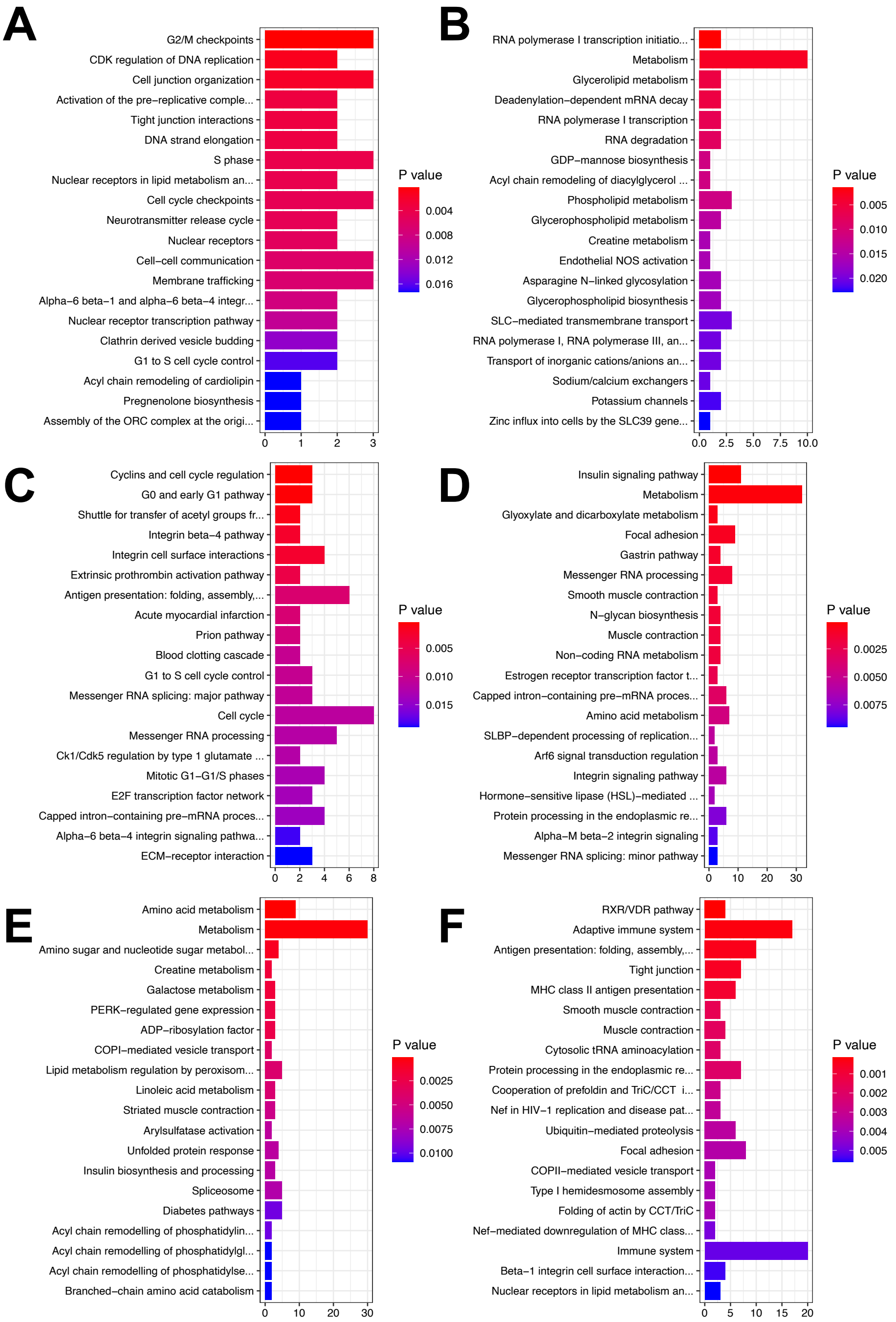


**Figure S1: Enrichr plots for differentially expressed genes at different developmental stages.** A) enriched pathways for downregulated and B) upregulated genes at 4hpf. C) enriched pathways for downregulated and D) upregulated genes at 6 hpf. E) enriched pathways for downregulated and F) upregulated genes at 8 hpf.

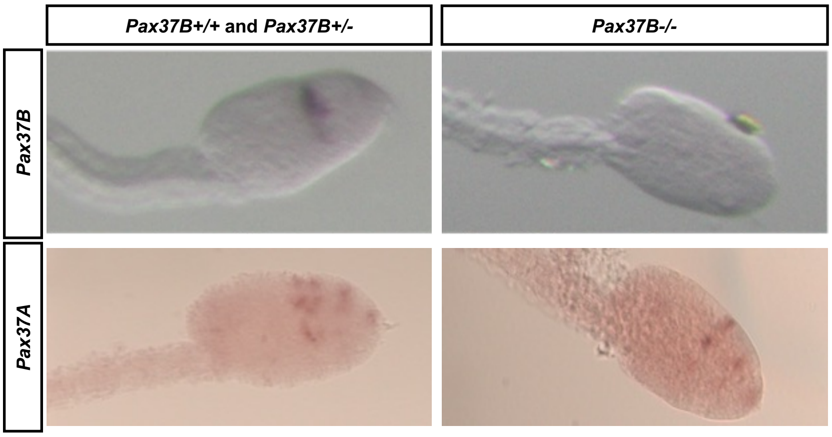


**Figure S2: ISH for Pax37B and Pax37A in embryos in crosses between *Pax37B$^{-/-}$* animals.** A) expression of *Pax37B* is not present in *Pax37B$^{-/-}$* animals suggesting a loss of Pax37B transcripts through nonsense-mediated decay. B) Expression of *Pax37A* seems to be unaffected in *Pax37B$^{-/-}$* animals compared to their heterozygote and wild-type siblings.

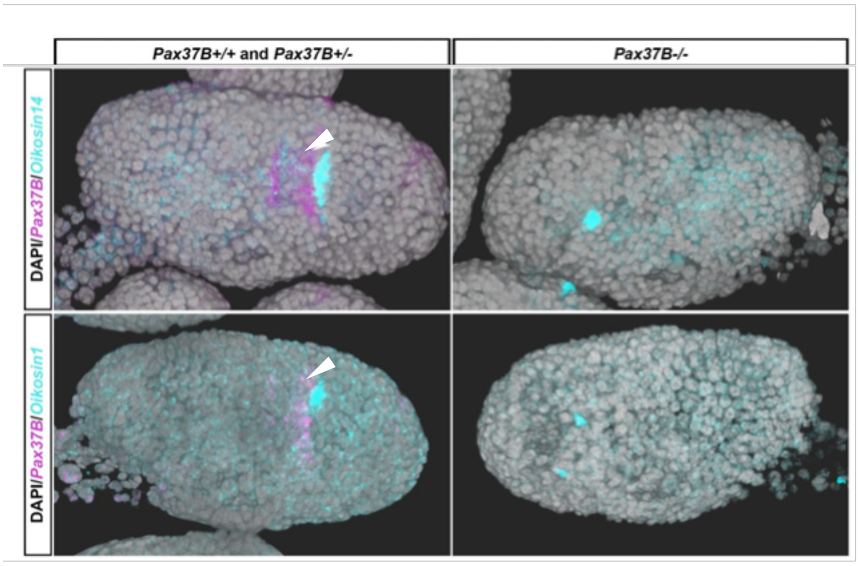


**Figure S3: FISH for differentially expressed giant Fol specific oikosins.** When performing FISH for oikosin1 and oikosin14 which are upregulated in the *Pax37B$^{-/-}$* animals we observed that some animals lacked staining and those that had, only were stained in lateral cell. White arrow point to *Pax37B* expression in Nasse cells.

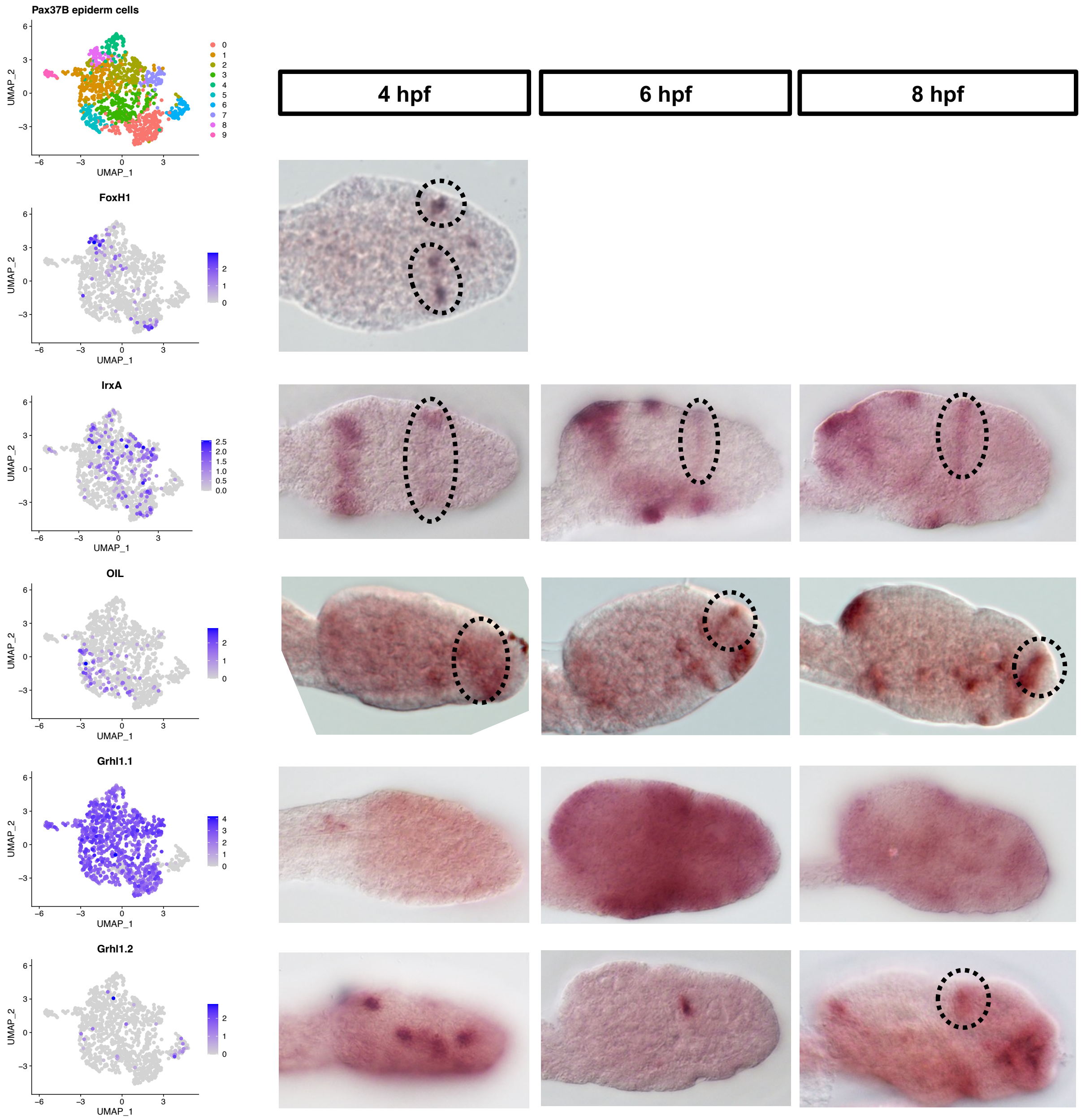


**Figure S4: Expression of different transcription factors that are lineage markers and/or differentially expressed in the *Pax37B+* epidermal subset and at three different developmental time-points.** The expression of these genes makes it possible to annotate the different clusters. *FoxH1* stops being expressed right after 4 hpf (Mikhaleva et al., 2018) and are thus only shown for one of the three stages. Dotted lines indicate epithelial expression that overlap with *Pax37B* expression.

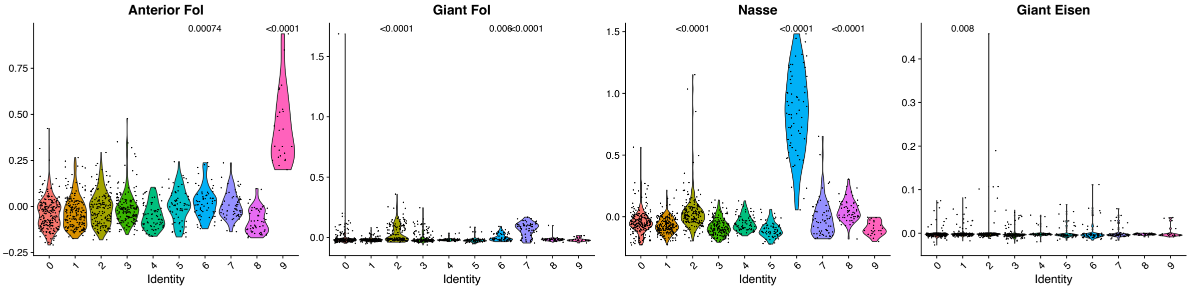


**Figure S5: Combined expression score of top 20 oikoplastic field specific marker genes in the *Pax37B+* epidermal cell subset reveal a higher expression of adult Nasse and adult giant Fol markers in clusters 2, 6, 7 and 8.** Statistics was calculated relative to basemean using Wilcoxon test with Bonferroni correction, only those with expression score above the basemean is shown.

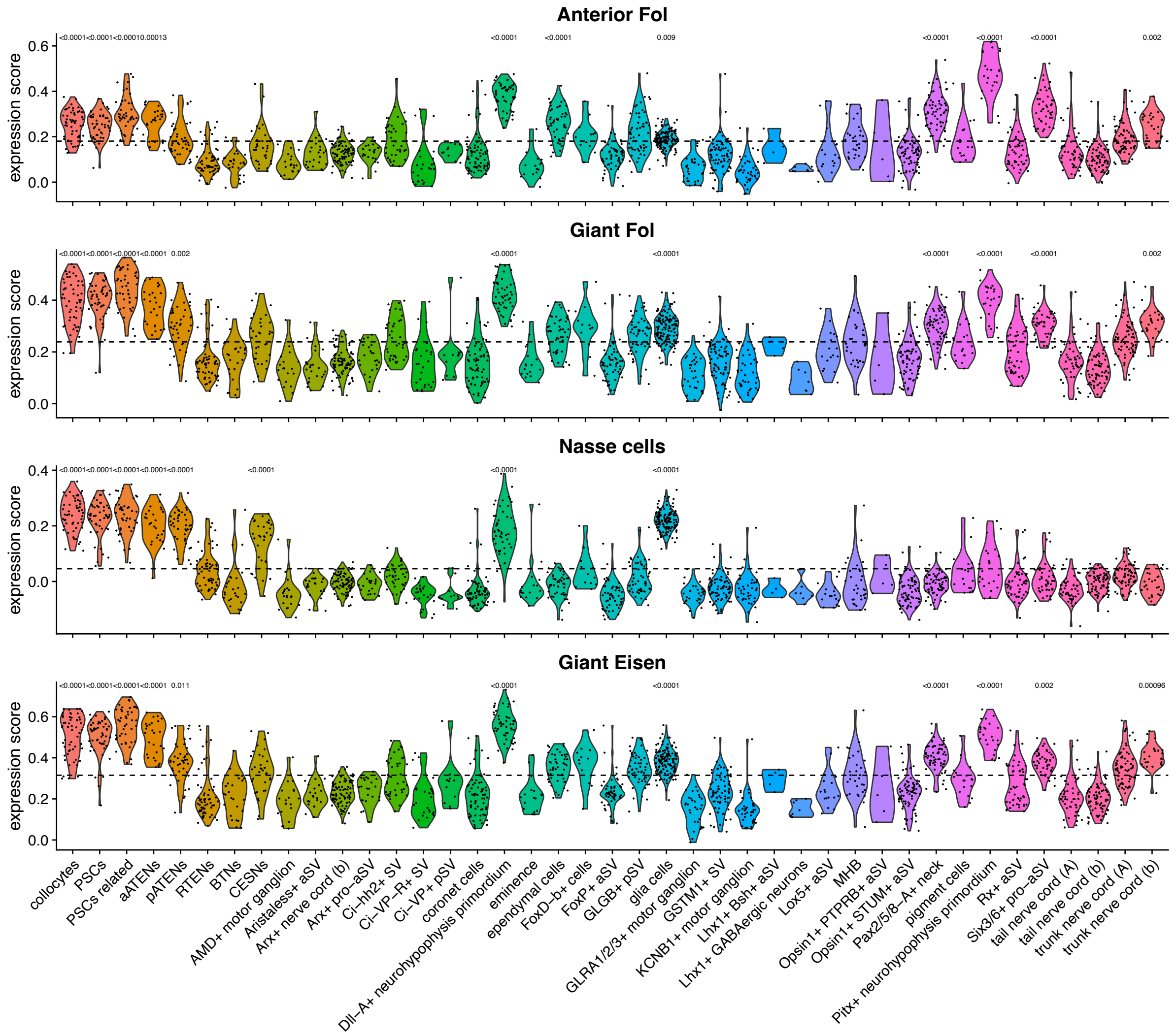


**Figure S6: Combined expression scores of Ciona robusta orthologs of oikoplastic field specific markers in Ciona robusta larval CNS.** Statistics was calculated relative to basemean using Wilcoxon test with Bonferroni correction, only those with expression score above the basemean is shown.
