## Supplementary material for "Pax37 gene function in *Oikopleura dioica* supports a neuroepithelial-like origin for its house-making Fol territory": R notebook with scripts used in this manuscript: Notebook_for_generating_analyses_and_plots_for_manuscript.nb.html

Scripts for analyses and plots presented in Functional analysis of O. diocia Pax37 genes in epiderm suggest a neuroepithelial-like origin of cells in the Fol region


Code 

- Show All Code
- Hide All Code
- Download Rmd

### Scripts for analyses and plots presented in Functional analysis of O. diocia Pax37 genes in epiderm suggest a neuroepithelial-like origin of cells in the Fol region

First a epiderm subset of our in house Oikopleura dioica scRNA-seq dataset was further subsetted into two objects, one with Pax37A and one with Pax37B cells. The following script is for generating the dataset and clustering for Pax37A positive epiderm cells:


```
# I set seed to ensure repeatability
set.seed(42)

##### SUBSET AND FIND MARKERS #####

# load the required packages
library(Seurat)
library(dplyr)
library(clustree)

# load the full dataset and normalize the data:
epiderm <- readRDS(file = "epi_2_0.6_clustering.rds")
DefaultAssay(epiderm) <- "RNA"
NormalizeData(epiderm)

# then subset the data based on expression of pax37a and output a file with the
# markers specific for pax37a expressing cells.
expression = GetAssayData(object = epiderm,
assay = "RNA", slot = "data")["GSOIDG00011822001:paired box gene 7",]

pos_ids = names(which(expression>0))
neg_ids = names(which(expression==0))
pos_cells = subset(epiderm,cells=pos_ids)

##### RECLUSTER SUBSET AND FIND MARKERS #####

DefaultAssay(pos_cells) <- "integrated"
pos_cells <- RunPCA(object = pos_cells,npcs = 100)

pdf(file="./pax37a/plots/PCA_elbow_epiderm.pdf", width = 10, height = 8)
ElbowPlot(pos_cells, ndims = 100, reduction = "pca")
dev.off()

pos_cells <- FindNeighbors(pos_cells, dims = 1:50)

# Select a range of resolutions
resolution.range <- seq(from = 0, to = 2, by = 0.2)

pos_cells <- Seurat::FindClusters(object = pos_cells, resolution = resolution.range)

pdf(file="./pax37a/plots/clustree_epiderm.pdf", width = 10, height = 8)
clustree(pos_cells)
dev.off()


### find markers for Pax37A positive epiderm cells ###
pax37a.full_de.markers <- FindMarkers(epiderm,
                                      ident.1 = pos_ids,
                                      ident.2 = neg_ids,
                                      only.pos = TRUE,
                                      min.pct = 0.25,
                                      logfc.threshold = 0.25)

sign.pax37a.full.de.markers <- pax37a.full_de.markers %>%
                                                      filter(p_val_adj < 0.05)

write.table(sign.pax37a.full.de.markers, file = "./pax37a/tables/sign.pax37a.epiderm.de.markers.txt",
            append = FALSE, sep = "\t", dec = ".",
            row.names = TRUE, col.names = TRUE)


##### RECLUSTER SUBSET AND FIND MARKERS #####
DefaultAssay(pos_cells) <- "integrated"
pos_cells <- RunPCA(object = pos_cells,npcs = 100)
pos_cells <- FindNeighbors(pos_cells, dims = 1:50)
pos_cells_1 <- Seurat::FindClusters(object = pos_cells, resolution = 2)
pos_cells_1 <- RunUMAP(pos_cells_1, dims = 1:50)

# print umaps of the reclusterings
pdf(file="./pax37a/plots/UMAP_epiderm.pdf", width = 10, height = 8)
DimPlot(pos_cells_1, reduction = "umap", label = TRUE)
dev.off()
pdf(file="./pax37a/plots/UMAP_orig.ident_epiderm.pdf", width = 10, height = 8)
DimPlot(pos_cells_1, reduction = "umap", group.by = "orig.ident")
dev.off()


# then make a markerlist for each cluster
DefaultAssay(pos_cells_1) <- "RNA"
pos_cells_1 <- NormalizeData(pos_cells_1)

pos_cells_1_epiderm.markers.Norm <- FindAllMarkers(pos_cells_1,
                                          assay = "RNA",
                                          slot = "data",
                                          only.pos = TRUE,
                                          min.pct = 0.25,
                                          logfc.threshold = 0.25)

sign.pos_cells_1_epiderm.markers.Norm <- pos_cells_1_epiderm.markers.Norm %>%
                                                          filter(p_val_adj < 0.05)

write.table(sign.pos_cells_1_epiderm.markers.Norm, file = "./pax37a/tables/sign.pos_cells_1.markers.Norm.txt",
            sep = '\t')

rm(epiderm)
save.image(file='./pax37a/pax37a_epiderm.RData')
```


The following script is to perform the same analysis but for Pax37B positive epidermal cells:


```
# I set seed to ensure repeatability
set.seed(42)

##### SUBSET AND FIND MARKERS #####

# load the required packages
library(Seurat)
library(dplyr)
library(clustree)

# load the full dataset and normalize the data:
epiderm <- readRDS(file = "epi_2_0.6_clustering.rds")
DefaultAssay(epiderm) <- "RNA"
NormalizeData(epiderm)

# then subset the data based on expression of Pax37B and output a file with the
# markers specific for Pax37B expressing cells.
expression = GetAssayData(object = epiderm,
assay = "RNA", slot = "data")["GSOIDG00011199001:paired box gene 7",]

pos_ids = names(which(expression>0))
neg_ids = names(which(expression==0))
pos_cells = subset(epiderm,cells=pos_ids)

##### RECLUSTER SUBSET AND FIND MARKERS #####

DefaultAssay(pos_cells) <- "integrated"
pos_cells <- RunPCA(object = pos_cells,npcs = 100)

pdf(file="./pax37b/plots/PCA_elbow_epiderm.pdf", width = 10, height = 8)
ElbowPlot(pos_cells, ndims = 100, reduction = "pca")
dev.off()

pos_cells <- FindNeighbors(pos_cells, dims = 1:50)

# Select a range of resolutions
resolution.range <- seq(from = 0, to = 2, by = 0.2)

pos_cells <- Seurat::FindClusters(object = pos_cells, resolution = resolution.range)

pdf(file="./pax37b/plots/clustree_epiderm.pdf", width = 10, height = 8)
clustree(pos_cells)
dev.off()


### find markers for Pax37B positive epidermal cells ###
pax37b.full_de.markers <- FindMarkers(epiderm,
                                      ident.1 = pos_ids,
                                      ident.2 = neg_ids,
                                      only.pos = TRUE,
                                      min.pct = 0.25,
                                      logfc.threshold = 0.25)

sign.pax37b.full.de.markers <- pax37b.full_de.markers %>%
                                                      filter(p_val_adj < 0.05)

write.table(sign.pax37b.full.de.markers, file = "./pax37b/tables/sign.pax37b.epiderm.de.markers.txt",
            append = FALSE, sep = "\t", dec = ".",
            row.names = TRUE, col.names = TRUE)


##### RECLUSTER SUBSET AND FIND MARKERS #####

DefaultAssay(pos_cells) <- "integrated"
pos_cells <- RunPCA(object = pos_cells,npcs = 100)
pos_cells <- FindNeighbors(pos_cells, dims = 1:50)
pos_cells_1 <- Seurat::FindClusters(object = pos_cells, resolution = 1.2)
pos_cells_1 <- RunUMAP(pos_cells_1, dims = 1:50)

# print umaps of the reclusterings
pdf(file="./pax37b/plots/UMAP_epiderm.pdf", width = 10, height = 8)
DimPlot(pos_cells_1, reduction = "umap", label = TRUE)
dev.off()
pdf(file="./pax37b/plots/UMAP_orig.ident_epiderm.pdf", width = 10, height = 8)
DimPlot(pos_cells_1, reduction = "umap", group.by = "orig.ident")
dev.off()


# then make a markerlist for each cluster
DefaultAssay(pos_cells_1) <- "RNA"
pos_cells_1 <- NormalizeData(pos_cells_1)

pos_cells_1_epiderm.markers.Norm <- FindAllMarkers(pos_cells_1,
                                          assay = "RNA",
                                          slot = "data",
                                          only.pos = TRUE,
                                          min.pct = 0.25,
                                          logfc.threshold = 0.25)

sign.pos_cells_1_epiderm.markers.Norm <- pos_cells_1_epiderm.markers.Norm %>%
                                                          filter(p_val_adj < 0.05)

write.table(sign.pos_cells_1_epiderm.markers.Norm, file = "./pax37b/tables/sign.pos_cells_1.markers.Norm.txt",
            sep = '\t')

rm(epiderm)

save.image(file='./pax37b/pax37b_epiderm.RData')
```


The next step is to generate the Pax37A split umap plot for Fig 2A in the manuscript:


```
library(Seurat)
library(dplyr)
library(ggplot2)
library(ggpubr)

# load the pax37a dataset
load("./pax37a/pax37a_epiderm.RData")

# Create a new metadata column called "stage" with the contents of "orig.ident" but with replicate number removed
pos_cells_1$stage <- gsub("[[:digit:]]", "", pos_cells_1$orig.ident)

# order stages
pos_cells_1$stage <- factor(x = pos_cells_1$stage, levels = c("four", "six", "eight","eleven","twelve","sixteen"))

# plot stages
stages <- DimPlot(pos_cells_1, label = FALSE, pt.size = 2, split.by = "stage") + labs(title = "Pax37A epiderm cells split by stage")
full <- DimPlot(pos_cells_1, label = FALSE, pt.size = 2) + xlim(-6,5) + ylim(-5,6) + labs(title = "Pax37A epiderm cells")

# generate plot of split umap
pdf(file="./pax37a/plots/split_by_stage_umap.pdf", width = 20, height = 5)
stages
dev.off()
```


Then the Pax37B split umap plot for Fig 2B in the manuscript:


```
library(Seurat)
library(dplyr)
library(ggplot2)
library(ggpubr)

# load the pax37b dataset
load("/Users/davidlagman/Desktop/Integrated_data_seurat/final_results/pax37b/pax37b_epiderm.RData")

# Create a new metadata column called "stage" with the contents of "orig.ident" but with replicate number removed
pos_cells_1$stage <- gsub("[[:digit:]]", "", pos_cells_1$orig.ident)

# order stages
pos_cells_1$stage <- factor(x = pos_cells_1$stage, levels = c("four", "six", "eight","eleven","twelve","sixteen"))

# plot stages
stages <- DimPlot(pos_cells_1, label = FALSE, pt.size = 2, split.by = "stage") + labs(title = "Pax37B epiderm cells split by stage")

# generate plot of split umap
pdf(file="./pax37b/plots/split_by_stage_umap.pdf", width = 20, height = 5)
stages
dev.off()
```


The follwing script is to perform the monocle3 analysis of Pax37B epidermal cells presented in Fig. 4A:


```
# I set seed to ensure repeatability
set.seed(42)

# load .RData file for pax37b
load(file='./pax37b/pax37b_epiderm.RData')

# load required libraries
library(Seurat)
library(SeuratWrappers)
library(monocle3)
library(dplyr)
library(viridis)
library(Polychrome)
library(ggplot2)

DefaultAssay(pos_cells_1) <-"RNA"

# Create a new metadata column called "stage" with the contents of "orig.ident" but with replicate number removed
pos_cells_1$stage <- gsub("[[:digit:]]", "", pos_cells_1$orig.ident)

monocle_object <- as.cell_data_set(pos_cells_1)
monocle_object <- cluster_cells(cds = monocle_object, reduction_method = "UMAP")
monocle_object <- learn_graph(monocle_object, use_partition = TRUE)

# a helper function to identify the root principal points:
get_earliest_principal_node <- function(monocle_object, time_bin="four"){
  cell_ids <- which(colData(monocle_object)[, "stage"] == time_bin)

  closest_vertex <-
    monocle_object@principal_graph_aux[["UMAP"]]$pr_graph_cell_proj_closest_vertex
  closest_vertex <- as.matrix(closest_vertex[colnames(monocle_object), ])
  root_pr_nodes <-
    igraph::V(principal_graph(monocle_object)[["UMAP"]])$name[as.numeric(names
                                                                     (which.max(table(closest_vertex[cell_ids,]))))]

  root_pr_nodes
}

monocle_object <- order_cells(monocle_object, root_pr_nodes=get_earliest_principal_node(monocle_object))

# plot trajectory
pdf(file="./pax37b/plots/monocle3_UMAP_epiderm.pdf", width = 10, height = 8)
plot_cells(
  cds = monocle_object,
  color_cells_by = "pseudotime",
  show_trajectory_graph = TRUE,
  cell_size = 2,
  trajectory_graph_segment_size = 2,
)
dev.off()

pos_cells_1 <- AddMetaData(
  object = pos_cells_1,
  metadata = monocle_object@principal_graph_aux@listData$UMAP$pseudotime,
  col.name = "pax37b"
)

# plot pseudotime on original umap
pdf(file="./pax37b/plots/pseudotime_UMAP_epiderm.pdf", width = 10, height = 8)
FeaturePlot(pos_cells_1, c("pax37b"), pt.size = 2) & scale_color_viridis_c()
dev.off()


# identify lineage markers for the principal lineage
pax37b_cds_pr_test_res <- graph_test(monocle_object, neighbor_graph="principal_graph", cores=8)
pr_deg_ids <- row.names(subset(pax37b_cds_pr_test_res, q_value < 1e-3))

write.table(pr_deg_ids, file = "./pax37b/tables/monocle3_analysis_epiderm_lineage_markers.txt")
```


The next step is to plot the Enrichr analysis presented in Fig. 4B and 4C.


```
library(enrichR)
library(ggpubr)

# script for making Enrichr plots for shared genes between Fol regions and de genes.

setEnrichrSite("Enrichr") # Human genes

websiteLive <- TRUE
dbs <- listEnrichrDbs()
if (is.null(dbs)) websiteLive <- FALSE
if (websiteLive) head(dbs)

down <- read.table("./pax37b/tables/annotated_lineage_markers_DOWN_reg_genes.txt")
up <- read.table("./pax37b/tables/annotated_lineage_markers_UP_reg_genes.txt")

dbs <- c("BioPlanet_2019")

if (websiteLive) {
  enriched_down <- enrichr(down$V1, dbs)
}

if (websiteLive) {
  enriched_up <- enrichr(up$V1, dbs)
}

downplot <- plotEnrich(enriched_down[[1]], showTerms = 20, numChar = 40, y = "Count", orderBy = "P.value") + theme(plot.title = element_blank(), axis.title.y = element_blank(), axis.title.x = element_blank())
upplot <- plotEnrich(enriched_up[[1]], showTerms = 20, numChar = 40, y = "Count", orderBy = "P.value") + theme(plot.title = element_blank(), axis.title.y = element_blank(), axis.title.x = element_blank())

pdf("./final_results/plots/enrichr_plot_shared_de_genes_lineage_markers.pdf", height = 4, width = 10)
ggarrange(downplot,upplot, ncol = 2, nrow = 1)
dev.off()
```


The next step is to plot the violin plots of the expression scores of down and up regulated oikosins in the Pax37B epidermal subset.


```
library(Seurat)
library(patchwork)
library(dplyr)
library(ggplot2)
# load the pax37b dataset
load("./pax37b/pax37b_epiderm.RData")
pax37b_up_oikosins <- read.table(file = "./pax37b/tables/upreg_putative_silix_oikosins.txt", sep = '\t', header = FALSE)
pax37b_down_oikosins <- read.table(file = "./pax37b/tables/downreg_putative_silix_oikosins.txt", sep = '\t', header = FALSE)

nasse <- pos_cells_1

pax37b_up_oikosins <- as.list(pax37b_up_oikosins$V1)
pax37b_down_oikosins <- as.list(pax37b_down_oikosins$V1)


nasse <- AddModuleScore(nasse,
                        features = list(intersect(rownames(nasse), pax37b_up_oikosins)),
                        name = "pax37b_up_oikosins")
nasse <- AddModuleScore(nasse,
                        features = list(intersect(rownames(nasse), pax37b_down_oikosins)),
                        name = "pax37b_down_oikosins")

pax37b_up_plot <- VlnPlot(nasse, features = "pax37b_up_oikosins1")

pax37b_down_plot <- VlnPlot(nasse, features = "pax37b_down_oikosins1")


pdf(file = "./pax37b/plots/de_oikosins_and_de_silix_oikosins_expresson_score.pdf", width = 15, height = 3.5)
pax37b_down_plot + ggtitle("Down oikosins") + 
  xlab("cluster") + 
  ylab("expression score") + 
  theme(legend.position = "none")  | pax37b_up_plot + 
  ggtitle("Up oikosins") + 
  xlab("cluster") + ylab("expression score") + 
  theme(legend.position = "none")
dev.off()
```


The next step is to generate the violin plots for the selection of putative developing Nasse and giant Fol clusters (Fig S5):


```
library(Seurat)
library(patchwork)
library(ggpubr)

# load the pax37b dataset
load("./pax37b/pax37b_epiderm.RData")

nasse_markers <- read.table("./nasse/tables/genenames_sign.nasse.de.markers.txt", sep = '\t', header = FALSE)
nasse_marker_gene_list <- head(nasse_markers, n = 20)

pos_cells_1 <- AddModuleScore(object = pos_cells_1, features = nasse_marker_gene_list, name = "nasse")

anteriorfol_markers <- read.table("./anterior_fol/tables/genenames_sign.anteriorfol.de.markers.txt", sep = '\t', header = FALSE)
anteriorfol_marker_gene_list <- head(anteriorfol_markers, n = 20)

pos_cells_1 <- AddModuleScore(object = pos_cells_1, features = anteriorfol_marker_gene_list, name = "anterior_fol")

giantfol_markers <- read.table("./giant_fol/tables/genenames_sign.giantfol.de.markers.txt", sep = '\t', header = FALSE)
giantfol_marker_gene_list <- head(giantfol_markers, n = 20)

pos_cells_1 <- AddModuleScore(object = pos_cells_1, features = giantfol_marker_gene_list, name = "giant_fol")


pos_cells_1 <- AddModuleScore(object = pos_cells_1, features = anteriorfol_marker_gene_list, name = "anterior_fol")

gianteisen_markers <- read.table("./giant_eisen/tables/genenames_sign.gianteisen.de.markers.txt", sep = '\t', header = FALSE)
gianteisen_marker_gene_list <- head(gianteisen_markers, n = 20)

pos_cells_1 <- AddModuleScore(object = pos_cells_1, features = gianteisen_marker_gene_list, name = "giant_eisen")


anteriorfolplot <- VlnPlot(pos_cells_1, features = "anterior_fol1") + labs(title = "Anterior Fol") + theme(legend.position = 'none') +geom_pwc(
  ref.group = "all", tip.length = 0,
  method = "wilcox_test", label = "p.adj.format",
  bracket.nudge.y = 0,
  p.adjust.method = "bonferroni",
  hide.ns = TRUE,
  label.size = 4,
  method.args = list(alternative = "less"))
nasseplot <- VlnPlot(pos_cells_1, features = "nasse1") + labs(title = "Nasse") + theme(legend.position = 'none')+geom_pwc(
  ref.group = "all", tip.length = 0,
  method = "wilcox_test", label = "p.adj.format",
  bracket.nudge.y = 0,
  p.adjust.method = "bonferroni",
  hide.ns = TRUE,
  label.size = 4,
  method.args = list(alternative = "less"))
giantfolplot <- VlnPlot(pos_cells_1, features = "giant_fol1") + labs(title = "Giant Fol") + theme(legend.position = 'none')+geom_pwc(
  ref.group = "all", tip.length = 0,
  method = "wilcox_test", label = "p.adj.format",
  bracket.nudge.y = 0,
  p.adjust.method = "bonferroni",
  hide.ns = TRUE,
  label.size = 4,
  method.args = list(alternative = "less"))
gianteisenplot <- VlnPlot(pos_cells_1, features = "giant_eisen1") + labs(title = "Giant Eisen") + theme(legend.position = 'none')+geom_pwc(
  ref.group = "all", tip.length = 0,
  method = "wilcox_test", label = "p.adj.format",
  bracket.nudge.y = 0,
  p.adjust.method = "bonferroni",
  hide.ns = TRUE,
  label.size = 4,
  method.args = list(alternative = "less"))

pdf(file="./pax37b/plots/top20_field_markers_expression_pax37bsubset.pdf", width = 20, height = 5)
anteriorfolplot | giantfolplot | nasseplot | gianteisenplot
dev.off()
```


The next step is to generate the putative developing Nasse and giant Fol subset and perform monocle3 lineage analysis and plot for Fig. 5A:


```
# I set seed to ensure repeatability
set.seed(42)

# load .RData file for pax37b
load(file='./pax37b/pax37b_epiderm.RData')

# load required libraries
library(Seurat)
library(clustree)
library(SeuratWrappers)
library(monocle3)
library(dplyr)
library(viridis)
library(Polychrome)
library(ggplot2)

# subset based on clusters identified in the previous step
nasse_lineage <- subset(pos_cells_1, idents = c("2","6","7","8"))
DimPlot(nasse_lineage)

##### RECLUSTER SUBSET AND FIND MARKERS #####

DefaultAssay(nasse_lineage) <- "integrated"
nasse_lineage <- RunPCA(object = nasse_lineage,npcs = 100)

pdf(file="./pax37b/plots/nasse_lineage_PCA_elbow_epiderm.pdf", width = 10, height = 8)
ElbowPlot(nasse_lineage, ndims = 100, reduction = "pca")
dev.off()

nasse_lineage <- FindNeighbors(nasse_lineage, dims = 1:25)

# Select a range of resolutions
resolution.range <- seq(from = 0, to = 2, by = 0.2)

nasse_lineage <- Seurat::FindClusters(object = nasse_lineage, resolution = resolution.range)

pdf(file="./pax37b/plots/nasse_lineage_clustree_epiderm.pdf", width = 10, height = 8)
clustree(nasse_lineage)
dev.off()


##### RECLUSTER SUBSET AND FIND MARKERS #####

DefaultAssay(nasse_lineage) <- "integrated"
nasse_lineage <- RunPCA(object = nasse_lineage,npcs = 100)
nasse_lineage <- FindNeighbors(nasse_lineage, dims = 1:25)
nasse_lineage_1 <- Seurat::FindClusters(object = nasse_lineage, resolution = 1)
nasse_lineage_1 <- RunUMAP(nasse_lineage_1, dims = 1:25)


DefaultAssay(nasse_lineage_1) <-"RNA"

# Create a new metadata column called "stage" with the contents of "orig.ident" but with replicate number removed
nasse_lineage_1$stage <- gsub("[[:digit:]]", "", nasse_lineage_1$orig.ident)

monocle_object <- as.cell_data_set(nasse_lineage_1)
monocle_object <- cluster_cells(cds = monocle_object, reduction_method = "UMAP")
monocle_object <- learn_graph(monocle_object, use_partition = TRUE)

# a helper function to identify the root principal points:
get_earliest_principal_node <- function(monocle_object, time_bin="four"){
  cell_ids <- which(colData(monocle_object)[, "stage"] == time_bin)
  
  closest_vertex <-
    monocle_object@principal_graph_aux[["UMAP"]]$pr_graph_cell_proj_closest_vertex
  closest_vertex <- as.matrix(closest_vertex[colnames(monocle_object), ])
  root_pr_nodes <-
    igraph::V(principal_graph(monocle_object)[["UMAP"]])$name[as.numeric(names
                                                                         (which.max(table(closest_vertex[cell_ids,]))))]
  
  root_pr_nodes
}

monocle_object <- order_cells(monocle_object, root_pr_nodes=get_earliest_principal_node(monocle_object))

# plot trajectory
pdf(file="./pax37b/plots/nasse_lineage_monocle3_UMAP_epiderm.pdf", width = 5, height = 5)
plot_cells(
  cds = monocle_object,
  color_cells_by = "pseudotime",
  show_trajectory_graph = TRUE,
  cell_size = 2,
  trajectory_graph_segment_size = 2
)
dev.off()

nasse_lineage <- AddMetaData(
  object = nasse_lineage_1,
  metadata = monocle_object@principal_graph_aux@listData$UMAP$pseudotime,
  col.name = "pax37b"
)

# identify lineage markers for the principal lineage
pax37b_cds_pr_test_res <- graph_test(monocle_object, neighbor_graph="principal_graph", cores=8)
pr_deg_ids <- row.names(subset(pax37b_cds_pr_test_res, q_value < 1e-3))

write.table(pr_deg_ids, file = "./pax37b/tables/nasse_lineage_monocle3_analysis_epiderm_lineage_markers.txt")
```


Next we create the heatmap over pseudotime for Fig 5B:


```
# I set seed to ensure repeatability
set.seed(42)

# load .RData file for pax37b
load(file='./pax37b/pax37b_epiderm.RData')

# load required libraries
library(Seurat)
library(SeuratWrappers)
library(monocle3)
library(dplyr)
library(ggplot2)
library(patchwork)
library(ComplexHeatmap)
library(RColorBrewer)
library(circlize)

nasse_lineage <- subset(pos_cells_1, idents = c("2","6","7","8"))

##### RECLUSTER SUBSET AND FIND MARKERS #####

DefaultAssay(nasse_lineage) <- "integrated"
nasse_lineage <- RunPCA(object = nasse_lineage,npcs = 100)
nasse_lineage <- FindNeighbors(nasse_lineage, dims = 1:25)
nasse_lineage_1 <- Seurat::FindClusters(object = nasse_lineage, resolution = 1)
nasse_lineage_1 <- RunUMAP(nasse_lineage_1, dims = 1:25)

DefaultAssay(nasse_lineage_1) <-"RNA"

nasse_lineage_1$stage <- gsub("[[:digit:]]", "", nasse_lineage_1$orig.ident)

monocle_object <- as.cell_data_set(nasse_lineage_1)
monocle_object <- cluster_cells(cds = monocle_object, reduction_method = "UMAP")
monocle_object <- learn_graph(monocle_object, use_partition = TRUE)

# a helper function to identify the root principal points:
get_earliest_principal_node <- function(monocle_object, time_bin="four"){
  cell_ids <- which(colData(monocle_object)[, "stage"] == time_bin)
  
  closest_vertex <-
    monocle_object@principal_graph_aux[["UMAP"]]$pr_graph_cell_proj_closest_vertex
  closest_vertex <- as.matrix(closest_vertex[colnames(monocle_object), ])
  root_pr_nodes <-
    igraph::V(principal_graph(monocle_object)[["UMAP"]])$name[as.numeric(names
                                                                         (which.max(table(closest_vertex[cell_ids,]))))]
  
  root_pr_nodes
}

monocle_object <- order_cells(monocle_object, root_pr_nodes=get_earliest_principal_node(monocle_object))

tfs <- read.table("./pax37b/tables/corrected_tfs_genenames_nasse_lineage_monocle3_analysis_epiderm_lineage_markers.txt", header = FALSE, stringsAsFactors = FALSE, sep = '\t')
tfs <- as.list(tfs)

pax37b_tfs <- tfs[["V1"]]

pt.matrix_tfs <- exprs(monocle_object)[match(pax37b_tfs,rownames(rowData(monocle_object))),order(pseudotime(monocle_object))]

pt.matrix_tfs <- t(apply(pt.matrix_tfs,1,function(x){smooth.spline(x,df=3)$y}))
pt.matrix_tfs <- t(apply(pt.matrix_tfs,1,function(x){(x-mean(x))/sd(x)}))
rownames(pt.matrix_tfs) <- pax37b_tfs;

rownames(pt.matrix_tfs) <- substr(rownames(pt.matrix_tfs),0,17)

df <- read.table("./pax37b/tables/for_dotplot_annotated_corrected_tfs_genenames_nasse_lineage_monocle3_analysis_epiderm_lineage_markers.txt", sep = "\t", header = FALSE, fill = TRUE)


pt.matrix_tfs <- cbind(rownames(pt.matrix_tfs), data.frame(pt.matrix_tfs, row.names=NULL))

indx <- match(pt.matrix_tfs$`rownames(pt.matrix_tfs)`, df$V1, nomatch = 0)
pt.matrix_tfs$`rownames(pt.matrix_tfs)`[indx != 0] <- df$V2[indx]

rownames(pt.matrix_tfs) <- pt.matrix_tfs$`rownames(pt.matrix_tfs)`

pt.matrix_tfs <- as.matrix(pt.matrix_tfs[,-1])


#Ward.D2 Hierarchical Clustering
hthc <- Heatmap(
  pt.matrix_tfs,
  name                         = "z-score",
  col                          = colorRamp2(seq(from=-2,to=2,length=11),rev(brewer.pal(11, "Spectral"))),
  show_row_names               = TRUE,
  show_column_names            = FALSE,
  row_names_gp                 = gpar(fontsize = 6),
  clustering_method_rows = "ward.D2",
  clustering_method_columns = "ward.D2",
  row_title_rot                = 0,
  cluster_rows                 = TRUE,
  cluster_row_slices           = FALSE,
  cluster_columns              = FALSE)

# print the heatmap and a FeaturePlot
pdf(file="./pax37b/plots/tfs_heatmap_nasse_lineage_markers.pdf", width = 5, height = 5)
print(hthc)
dev.off()
```


The next step is to generate the clustered dotplots for the expression of Ciona robusta orthologs of Nasse and putative giant Fol precursor lineage transcription factors for Fig. 9A and 9B.


```
# set maximum ram usage:
options(future.globals.maxSize = 64000 * 1024^2)

# I set seed to ensure repeatability
set.seed(42)

library(circlize)
library(ComplexHeatmap)
library(cowplot)
library(data.table)
library(ggplot2)
library(patchwork)
library(plyr)
library(Polychrome)
library(reshape2)
library(Seurat)
library(tidyverse)
library(UpSetR)
library(viridis)
library(clustree)


expr <- fread("ciona/expression_matrix_10stage.tsv", header = TRUE, row.names(1))
meta <- fread("edited_ciona10stage.cluster.upload.new.txt", header = TRUE)

rownames(meta) <- meta$NAME
rownames(expr) <- expr$GENE
expr <- as.data.frame(expr)
expr2 <- expr[,-1]
rownames(expr2) <- expr$GENE
full <- CreateSeuratObject(counts = expr2, meta.data = meta)

# subset epidermal cells
epiderm <- subset(full, subset = Tissue.Type == "epidermis")

# subset epidermal cells that express pax37
DefaultAssay(epiderm) <- "RNA"
NormalizeData(epiderm)

# then subset the data based on expression of Pax37B and output a file with the
# markers specific for Pax37B expressing cells.
expression = GetAssayData(object = epiderm,
                          assay = "RNA", slot = "data")["KH2012:KH.C10.150",]

pos_ids = names(which(expression>0))
neg_ids = names(which(expression==0))
pos_cells = subset(epiderm,cells=pos_ids)

# Create a new metadata column called "stage" with the contents of "orig.ident" but with replicate number removed
pos_cells$orig.ident <- substr(pos_cells$orig.ident,1,4)

pos_cells$orig.ident <- gsub("lv.1", "larvae", pos_cells$orig.ident)
pos_cells$orig.ident <- gsub("lv.3", "larvae", pos_cells$orig.ident)
pos_cells$orig.ident <- gsub("lv.4", "larvae", pos_cells$orig.ident)

pos_cells$orig.ident <- gsub("midG", "mid gastrula", pos_cells$orig.ident)
pos_cells$orig.ident <- gsub("earl", "early neurula", pos_cells$orig.ident)
pos_cells$orig.ident <- gsub("late", "late neurula", pos_cells$orig.ident)
pos_cells$orig.ident <- gsub("ITB.", "initial tailbud", pos_cells$orig.ident)
pos_cells$orig.ident <- gsub("ETB.", "early tailbud", pos_cells$orig.ident)
pos_cells$orig.ident <- gsub("MTB.", "mid tailbud", pos_cells$orig.ident)
pos_cells$orig.ident <- gsub("LTB1", "late tailbud I", pos_cells$orig.ident)
pos_cells$orig.ident <- gsub("LTB2", "late tailbud II", pos_cells$orig.ident)

orthlin <- fread("./pax37b/tables/ciona_orthologs_corrected_tfs_genenames_nasse_lineage_monocle3_analysis_epiderm_lineage_markers.txt", header = FALSE)

dp <- DotPlot(pos_cells, features = orthlin$V1, group.by = "orig.ident")

ddf<- dp$data

### the matrix for the scaled expression 
dexp_mat <- ddf %>% 
  select(-pct.exp, -avg.exp) %>%  
  pivot_wider(names_from = id, values_from = avg.exp.scaled) %>% 
  as.data.frame() 

dpercent_mat<-ddf %>% 
  select(-avg.exp, -avg.exp.scaled) %>%  
  pivot_wider(names_from = id, values_from = pct.exp) %>% 
  as.data.frame() 

dexp_mat2 <- dexp_mat[,-1]
rownames(dexp_mat2) <- dexp_mat$features.plot
dpercent_mat2 <- dpercent_mat[,-1]
rownames(dpercent_mat2) <- dpercent_mat$features.plot

dexp <- as.matrix(sapply(dexp_mat2, as.numeric))  
rownames(dexp) <-  dexp_mat$features.plot
dperc <- as.matrix(sapply(dpercent_mat2, as.numeric))
rownames(dperc) <-  dpercent_mat$features.plot

## any value that is greater than 2 will be mapped to yellow
dcol_fun = circlize::colorRamp2(c(-2, 0, 2), viridis(20)[c(1,10, 20)])

dcell_fun = function(j, i, x, y, w, h, fill){
  grid.rect(x = x, y = y, width = w, height = h, 
            gp = gpar(col = NA, fill = NA))
  grid.circle(x=x,y=y,r= dperc[i, j]/100 * min(unit.c(w, h)),
              gp = gpar(fill = dcol_fun(dexp[i, j]), col = NA))}

df <- read.table("./pax37b/tables/with_ciona_orthologs_and_annotations_corrected_tfs_genenames_nasse_lineage_monocle3_analysis_epiderm_lineage_markers.txt", sep = "\t", header = FALSE, fill = TRUE)


dexp <- cbind(rownames(dexp), data.frame(dexp, row.names=NULL))

indx <- match(dexp$`rownames(dexp)`, df$V3, nomatch = 0)
dexp$`rownames(dexp)`[indx != 0] <- df$V2[indx]

rownames(dexp) <- dexp$`rownames(dexp)`

dexp <- as.matrix(dexp[,-1])

map <- Heatmap(dexp,
                  heatmap_legend_param=list(title="avg expression", legend_direction = "vertical"),
                  column_title = "Nasse lineage TFs ciona epiderm", 
                  col=dcol_fun,
                  rect_gp = gpar(type = "none"),
                  cell_fun = dcell_fun,
                  row_names_gp = gpar(fontsize = 5),
                  column_names_gp = gpar(fontsize = 5),
                  border = "black",
                  column_names_rot = 90)

pdf("./pax37b/plots/expression_of_nasse_lineage_TFs_ciona_epiderm.pdf", width = 4, height = 4)
map
dev.off()

### Nervous system ###

# subset neuronal cells
neuron <- subset(full, subset = Tissue.Type == "nervous system")

# subset neuronal cells that express pax37
DefaultAssay(neuron) <- "RNA"
NormalizeData(neuron)

# then subset the data based on expression of Pax37B and output a file with the
# markers specific for Pax37B expressing cells.
expression = GetAssayData(object = neuron,
                          assay = "RNA", slot = "data")["KH2012:KH.C10.150",]

pos_ids = names(which(expression>0))
neg_ids = names(which(expression==0))
pos_cells = subset(neuron,cells=pos_ids)

# Create a new metadata column called "stage" with the contents of "orig.ident" but with replicate number removed
pos_cells$orig.ident <- substr(pos_cells$orig.ident,1,4)

pos_cells$orig.ident <- gsub("lv.1", "larvae", pos_cells$orig.ident)
pos_cells$orig.ident <- gsub("lv.3", "larvae", pos_cells$orig.ident)
pos_cells$orig.ident <- gsub("lv.4", "larvae", pos_cells$orig.ident)

pos_cells$orig.ident <- gsub("midG", "mid gastrula", pos_cells$orig.ident)
pos_cells$orig.ident <- gsub("earl", "early neurula", pos_cells$orig.ident)
pos_cells$orig.ident <- gsub("late", "late neurula", pos_cells$orig.ident)
pos_cells$orig.ident <- gsub("ITB.", "initial tailbud", pos_cells$orig.ident)
pos_cells$orig.ident <- gsub("ETB.", "early tailbud", pos_cells$orig.ident)
pos_cells$orig.ident <- gsub("MTB.", "mid tailbud", pos_cells$orig.ident)
pos_cells$orig.ident <- gsub("LTB1", "late tailbud I", pos_cells$orig.ident)
pos_cells$orig.ident <- gsub("LTB2", "late tailbud II", pos_cells$orig.ident)

orthlin <- fread("./pax37b/tables/ciona_orthologs_corrected_tfs_genenames_nasse_lineage_monocle3_analysis_epiderm_lineage_markers.txt", header = FALSE)

dp <- DotPlot(pos_cells, features = orthlin$V1, group.by = "orig.ident")

ddf<- dp$data

### the matrix for the scaled expression 
dexp_mat <- ddf %>% 
  select(-pct.exp, -avg.exp) %>%  
  pivot_wider(names_from = id, values_from = avg.exp.scaled) %>% 
  as.data.frame() 

dpercent_mat<-ddf %>% 
  select(-avg.exp, -avg.exp.scaled) %>%  
  pivot_wider(names_from = id, values_from = pct.exp) %>% 
  as.data.frame() 

dexp_mat2 <- dexp_mat[,-1]
rownames(dexp_mat2) <- dexp_mat$features.plot
dpercent_mat2 <- dpercent_mat[,-1]
rownames(dpercent_mat2) <- dpercent_mat$features.plot

dexp <- as.matrix(sapply(dexp_mat2, as.numeric))  
rownames(dexp) <-  dexp_mat$features.plot
dperc <- as.matrix(sapply(dpercent_mat2, as.numeric))
rownames(dperc) <-  dpercent_mat$features.plot

## any value that is greater than 2 will be mapped to yellow
dcol_fun = circlize::colorRamp2(c(-2, 0, 2), viridis(20)[c(1,10, 20)])

dcell_fun = function(j, i, x, y, w, h, fill){
  grid.rect(x = x, y = y, width = w, height = h, 
            gp = gpar(col = NA, fill = NA))
  grid.circle(x=x,y=y,r= dperc[i, j]/100 * min(unit.c(w, h)),
              gp = gpar(fill = dcol_fun(dexp[i, j]), col = NA))}

df <- read.table("./pax37b/tables/with_ciona_orthologs_and_annotations_corrected_tfs_genenames_nasse_lineage_monocle3_analysis_epiderm_lineage_markers.txt", sep = "\t", header = FALSE, fill = TRUE)


dexp <- cbind(rownames(dexp), data.frame(dexp, row.names=NULL))

indx <- match(dexp$`rownames(dexp)`, df$V3, nomatch = 0)
dexp$`rownames(dexp)`[indx != 0] <- df$V2[indx]

rownames(dexp) <- dexp$`rownames(dexp)`

dexp <- as.matrix(dexp[,-1])

map <- Heatmap(dexp,
                  heatmap_legend_param=list(title="avg expression", legend_direction = "vertical"),
                  column_title = "Nasse lineage TFs ciona nervous system", 
                  col=dcol_fun,
                  rect_gp = gpar(type = "none"),
                  cell_fun = dcell_fun,
                  row_names_gp = gpar(fontsize = 5),
                  column_names_gp = gpar(fontsize = 5),
                  border = "black",
                  column_names_rot = 90)

pdf("./pax37b/plots/expression_of_nasse_lineage_TFs_ciona_nervous_system.pdf", width = 4, height = 4)
map
dev.off()
```


Next we generate the heatmaps of scaled expression scores of orthologs to O. dioica cluster markers in the full and CNS Ciona robusta dataset for Fig. 10:


```
library(Seurat)
library(data.table)
library(patchwork)
library(ggpubr)
library(cowplot)
library(ggplot2)
library(plyr)
library(reshape2)
library(tidyverse)
library(dplyr)
library(stringr)
library(pheatmap)
library(gridExtra)

# load the larval cns dataset
expr <- fread("CNS.lv.expressionmatrix.rename.tsv", header = TRUE, row.names(1))
meta <- fread("CNS.lv.clusters.upload.renameedit.txt", header = TRUE)

rownames(meta) <- meta$NAME
rownames(expr) <- expr$GENE
expr <- as.data.frame(expr)
expr2 <- expr[,-1]
rownames(expr2) <- expr$GENE
larvalcns <- CreateSeuratObject(counts = expr2, meta.data = meta)

# load the full dataset
exprfull <- fread("expression_matrix_10stage.tsv", header = TRUE, row.names(1))
metafull <- fread("edited_ciona10stage.cluster.upload.new.txt", header = TRUE)

rownames(metafull) <- metafull$NAME
rownames(exprfull) <- exprfull$GENE
exprfull <- as.data.frame(exprfull)
exprfull2 <- exprfull[,-1]
rownames(exprfull2) <- exprfull$GENE
full <- CreateSeuratObject(counts = exprfull2, meta.data = metafull)

pax37b0 <- fread("./pax37b/tables/cluster_markers/ciona_orthologs_0_markers.txt", header = FALSE, sep = '\t')
pax37b1 <- fread("./pax37b/tables/cluster_markers/ciona_orthologs_1_markers.txt", header = FALSE, sep = '\t')
pax37b2 <- fread("./pax37b/tables/cluster_markers/ciona_orthologs_2_markers.txt", header = FALSE, sep = '\t')
pax37b3 <- fread("./pax37b/tables/cluster_markers/ciona_orthologs_3_markers.txt", header = FALSE, sep = '\t')
pax37b4 <- fread("./pax37b/tables/cluster_markers/ciona_orthologs_4_markers.txt", header = FALSE, sep = '\t')
pax37b5 <- fread("./pax37b/tables/cluster_markers/ciona_orthologs_5_markers.txt", header = FALSE, sep = '\t')
pax37b6 <- fread("./pax37b/tables/cluster_markers/ciona_orthologs_6_markers.txt", header = FALSE, sep = '\t')
pax37b7 <- fread("./pax37b/tables/cluster_markers/ciona_orthologs_7_markers.txt", header = FALSE, sep = '\t')
pax37b8 <- fread("./pax37b/tables/cluster_markers/ciona_orthologs_8_markers.txt", header = FALSE, sep = '\t')
pax37b9 <- fread("./pax37b/tables/cluster_markers/ciona_orthologs_9_markers.txt", header = FALSE, sep = '\t')

cns0<- fread("./cns/cluster_markers/only_ciona_orthologs_0_markers.txt", header = FALSE, sep = '\t')
cns1<- fread("./cns/cluster_markers/only_ciona_orthologs_1_markers.txt", header = FALSE, sep = '\t')
cns2<- fread("./cns/cluster_markers/only_ciona_orthologs_2_markers.txt", header = FALSE, sep = '\t')
cns3<- fread("./cns/cluster_markers/only_ciona_orthologs_3_markers.txt", header = FALSE, sep = '\t')
cns4<- fread("./cns/cluster_markers/only_ciona_orthologs_4_markers.txt", header = FALSE, sep = '\t')
cns5<- fread("./cns/cluster_markers/only_ciona_orthologs_5_markers.txt", header = FALSE, sep = '\t')
cns6<- fread("./cns/cluster_markers/only_ciona_orthologs_6_markers.txt", header = FALSE, sep = '\t')
cns7<- fread("./cns/cluster_markers/only_ciona_orthologs_7_markers.txt", header = FALSE, sep = '\t')
cns8<- fread("./cns/cluster_markers/only_ciona_orthologs_8_markers.txt", header = FALSE, sep = '\t')
cns9<- fread("./cns/cluster_markers/only_ciona_orthologs_9_markers.txt", header = FALSE, sep = '\t')
cns10<- fread("./cns/cluster_markers/only_ciona_orthologs_10_markers.txt", header = FALSE, sep = '\t')
cns11<- fread("./cns/cluster_markers/only_ciona_orthologs_11_markers.txt", header = FALSE, sep = '\t')
cns12<- fread("./cns/cluster_markers/only_ciona_orthologs_12_markers.txt", header = FALSE, sep = '\t')
cns13<- fread("./cns/cluster_markers/only_ciona_orthologs_13_markers.txt", header = FALSE, sep = '\t')
cns14<- fread("./cns/cluster_markers/only_ciona_orthologs_14_markers.txt", header = FALSE, sep = '\t')
cns15<- fread("./cns/cluster_markers/only_ciona_orthologs_15_markers.txt", header = FALSE, sep = '\t')
cns16<- fread("./cns/cluster_markers/only_ciona_orthologs_16_markers.txt", header = FALSE, sep = '\t')
cns17<- fread("./cns/cluster_markers/only_ciona_orthologs_17_markers.txt", header = FALSE, sep = '\t')
cns18<- fread("./cns/cluster_markers/only_ciona_orthologs_18_markers.txt", header = FALSE, sep = '\t')
cns19<- fread("./cns/cluster_markers/only_ciona_orthologs_19_markers.txt", header = FALSE, sep = '\t')
cns20<- fread("./cns/cluster_markers/only_ciona_orthologs_20_markers.txt", header = FALSE, sep = '\t')

pax37b396_4hpf_down <- fread("./de_genes_pax37b396_mutants/geneids/ciona_orthologs_4_hpf_DOWN_geneids_20200317.txt", header = FALSE, sep = '\t')
pax37b396_4hpf_up <- fread("./de_genes_pax37b396_mutants/geneids/ciona_orthologs_4_hpf_UP_geneids_20200317.txt", header = FALSE, sep = '\t')
pax37b396_6hpf_down <- fread("./de_genes_pax37b396_mutants/geneids/ciona_orthologs_6hpf_DOWN_geneids_20200303.txt", header = FALSE, sep = '\t')
pax37b396_6hpf_up <- fread("./de_genes_pax37b396_mutants/geneids/ciona_orthologs_6hpf_UP_geneids_20200303.txt", header = FALSE, sep = '\t')
pax37b396_8hpf_down <- fread("./de_genes_pax37b396_mutants/geneids/ciona_orthologs_8hpf_DOWN_geneids_20200303.txt", header = FALSE, sep = '\t')
pax37b396_8hpf_up <- fread("./de_genes_pax37b396_mutants/geneids/ciona_orthologs_8hpf_UP_geneids_20200303.txt", header = FALSE, sep = '\t')


larvalcns <- AddModuleScore(larvalcns,
                            features = list(intersect(rownames(larvalcns), pax37b396_4hpf_down$V1)),
                            name = "4_DOWN")
larvalcns <- AddModuleScore(larvalcns,
                            features = list(intersect(rownames(larvalcns), pax37b396_4hpf_up$V1)),
                            name = "4_UP")
larvalcns <- AddModuleScore(larvalcns,
                            features = list(intersect(rownames(larvalcns), pax37b396_6hpf_down$V1)),
                            name = "6_DOWN")
larvalcns <- AddModuleScore(larvalcns,
                            features = list(intersect(rownames(larvalcns), pax37b396_6hpf_up$V1)),
                            name = "6_UP")
larvalcns <- AddModuleScore(larvalcns,
                            features = list(intersect(rownames(larvalcns), pax37b396_8hpf_down$V1)),
                            name = "8_DOWN")
larvalcns <- AddModuleScore(larvalcns,
                            features = list(intersect(rownames(larvalcns), pax37b396_8hpf_up$V1)),
                            name = "8_UP")


larvalcns <- AddModuleScore(larvalcns,
                            features = list(intersect(rownames(larvalcns), pax37b0$V1)),
                            name = "pax37b0")
larvalcns <- AddModuleScore(larvalcns,
                            features = list(intersect(rownames(larvalcns), pax37b1$V1)),
                            name = "pax37b1")
larvalcns <- AddModuleScore(larvalcns,
                            features = list(intersect(rownames(larvalcns), pax37b2$V1)),
                            name = "pax37b2")
larvalcns <- AddModuleScore(larvalcns,
                            features = list(intersect(rownames(larvalcns), pax37b3$V1)),
                            name = "pax37b3")
larvalcns <- AddModuleScore(larvalcns,
                            features = list(intersect(rownames(larvalcns), pax37b4$V1)),
                            name = "pax37b4")
larvalcns <- AddModuleScore(larvalcns,
                            features = list(intersect(rownames(larvalcns), pax37b5$V1)),
                            name = "pax37b5")
larvalcns <- AddModuleScore(larvalcns,
                            features = list(intersect(rownames(larvalcns), pax37b6$V1)),
                            name = "pax37b6")
larvalcns <- AddModuleScore(larvalcns,
                            features = list(intersect(rownames(larvalcns), pax37b7$V1)),
                            name = "pax37b7")
larvalcns <- AddModuleScore(larvalcns,
                            features = list(intersect(rownames(larvalcns), pax37b8$V1)),
                            name = "pax37b8")
larvalcns <- AddModuleScore(larvalcns,
                            features = list(intersect(rownames(larvalcns), pax37b9$V1)),
                            name = "pax37b9")

larvalcns <- AddModuleScore(larvalcns,      
                            features = list(intersect(rownames(larvalcns), cns0$V1)),
                            name = "cns0")
larvalcns <- AddModuleScore(larvalcns,      
                            features = list(intersect(rownames(larvalcns), cns1$V1)),
                            name = "cns1")
larvalcns <- AddModuleScore(larvalcns,      
                            features = list(intersect(rownames(larvalcns), cns2$V1)),
                            name = "cns2")
larvalcns <- AddModuleScore(larvalcns,      
                            features = list(intersect(rownames(larvalcns), cns3$V1)),
                            name = "cns3")
larvalcns <- AddModuleScore(larvalcns,      
                            features = list(intersect(rownames(larvalcns), cns4$V1)),
                            name = "cns4")
larvalcns <- AddModuleScore(larvalcns,      
                            features = list(intersect(rownames(larvalcns), cns5$V1)),
                            name = "cns5")
larvalcns <- AddModuleScore(larvalcns,      
                            features = list(intersect(rownames(larvalcns), cns6$V1)),
                            name = "cns6")
larvalcns <- AddModuleScore(larvalcns,      
                            features = list(intersect(rownames(larvalcns),cns7$V1)),
                            name = "cns7")
larvalcns <- AddModuleScore(larvalcns,      
                            features = list(intersect(rownames(larvalcns), cns8$V1)),
                            name = "cns8")
larvalcns <- AddModuleScore(larvalcns,      
                            features = list(intersect(rownames(larvalcns), cns9$V1)),
                            name = "cns9")
larvalcns <- AddModuleScore(larvalcns,      
                            features = list(intersect(rownames(larvalcns), cns10$V1)),
                            name = "cns10")
larvalcns <- AddModuleScore(larvalcns,      
                            features = list(intersect(rownames(larvalcns), cns11$V1)),
                            name = "cns11")
larvalcns <- AddModuleScore(larvalcns,      
                            features = list(intersect(rownames(larvalcns), cns12$V1)),
                            name = "cns12")
larvalcns <- AddModuleScore(larvalcns,      
                            features = list(intersect(rownames(larvalcns), cns13$V1)),
                            name = "cns13")
larvalcns <- AddModuleScore(larvalcns,      
                            features = list(intersect(rownames(larvalcns),cns14$V1)),
                            name = "cns14")
larvalcns <- AddModuleScore(larvalcns,      
                            features = list(intersect(rownames(larvalcns), cns15$V1)),
                            name = "cns15")
larvalcns <- AddModuleScore(larvalcns,      
                            features = list(intersect(rownames(larvalcns), cns16$V1)),
                            name = "cns16")
larvalcns <- AddModuleScore(larvalcns,      
                            features = list(intersect(rownames(larvalcns), cns17$V1)),
                            name = "cns17")
larvalcns <- AddModuleScore(larvalcns,      
                            features = list(intersect(rownames(larvalcns), cns18$V1)),
                            name = "cns18")
larvalcns <- AddModuleScore(larvalcns,      
                            features = list(intersect(rownames(larvalcns), cns19$V1)),
                            name = "cns19")
larvalcns <- AddModuleScore(larvalcns,      
                            features = list(intersect(rownames(larvalcns), cns20$V1)),
                            name = "cns20")

full <- AddModuleScore(full,
                            features = list(intersect(rownames(full), pax37b396_4hpf_down$V1)),
                            name = "4_DOWN")
full <- AddModuleScore(full,
                            features = list(intersect(rownames(full), pax37b396_4hpf_up$V1)),
                            name = "4_UP")
full <- AddModuleScore(full,
                            features = list(intersect(rownames(full), pax37b396_6hpf_down$V1)),
                            name = "6_DOWN")
full <- AddModuleScore(full,
                            features = list(intersect(rownames(full), pax37b396_6hpf_up$V1)),
                            name = "6_UP")
full <- AddModuleScore(full,
                            features = list(intersect(rownames(full), pax37b396_8hpf_down$V1)),
                            name = "8_DOWN")
full <- AddModuleScore(full,
                            features = list(intersect(rownames(full), pax37b396_8hpf_up$V1)),
                            name = "8_UP")


full <- AddModuleScore(full,
                            features = list(intersect(rownames(full), pax37b0$V1)),
                            name = "pax37b0")
full <- AddModuleScore(full,
                            features = list(intersect(rownames(full), pax37b1$V1)),
                            name = "pax37b1")
full <- AddModuleScore(full,
                            features = list(intersect(rownames(full), pax37b2$V1)),
                            name = "pax37b2")
full <- AddModuleScore(full,
                            features = list(intersect(rownames(full), pax37b3$V1)),
                            name = "pax37b3")
full <- AddModuleScore(full,
                            features = list(intersect(rownames(full), pax37b4$V1)),
                            name = "pax37b4")
full <- AddModuleScore(full,
                            features = list(intersect(rownames(full), pax37b5$V1)),
                            name = "pax37b5")
full <- AddModuleScore(full,
                            features = list(intersect(rownames(full), pax37b6$V1)),
                            name = "pax37b6")
full <- AddModuleScore(full,
                            features = list(intersect(rownames(full), pax37b7$V1)),
                            name = "pax37b7")
full <- AddModuleScore(full,
                            features = list(intersect(rownames(full), pax37b8$V1)),
                            name = "pax37b8")
full <- AddModuleScore(full,
                            features = list(intersect(rownames(full), pax37b9$V1)),
                            name = "pax37b9")

full <- AddModuleScore(full,        
                            features = list(intersect(rownames(full), cns0$V1)),
                            name = "cns0")
full <- AddModuleScore(full,        
                            features = list(intersect(rownames(full), cns1$V1)),
                            name = "cns1")
full <- AddModuleScore(full,        
                            features = list(intersect(rownames(full), cns2$V1)),
                            name = "cns2")
full <- AddModuleScore(full,        
                            features = list(intersect(rownames(full), cns3$V1)),
                            name = "cns3")
full <- AddModuleScore(full,        
                            features = list(intersect(rownames(full), cns4$V1)),
                            name = "cns4")
full <- AddModuleScore(full,        
                            features = list(intersect(rownames(full), cns5$V1)),
                            name = "cns5")
full <- AddModuleScore(full,        
                            features = list(intersect(rownames(full), cns6$V1)),
                            name = "cns6")
full <- AddModuleScore(full,        
                            features = list(intersect(rownames(full),cns7$V1)),
                            name = "cns7")
full <- AddModuleScore(full,        
                            features = list(intersect(rownames(full), cns8$V1)),
                            name = "cns8")
full <- AddModuleScore(full,        
                            features = list(intersect(rownames(full), cns9$V1)),
                            name = "cns9")
full <- AddModuleScore(full,        
                            features = list(intersect(rownames(full), cns10$V1)),
                            name = "cns10")
full <- AddModuleScore(full,        
                            features = list(intersect(rownames(full), cns11$V1)),
                            name = "cns11")
full <- AddModuleScore(full,        
                            features = list(intersect(rownames(full), cns12$V1)),
                            name = "cns12")
full <- AddModuleScore(full,        
                            features = list(intersect(rownames(full), cns13$V1)),
                            name = "cns13")
full <- AddModuleScore(full,        
                            features = list(intersect(rownames(full),cns14$V1)),
                            name = "cns14")
full <- AddModuleScore(full,        
                            features = list(intersect(rownames(full), cns15$V1)),
                            name = "cns15")
full <- AddModuleScore(full,        
                            features = list(intersect(rownames(full), cns16$V1)),
                            name = "cns16")
full <- AddModuleScore(full,        
                            features = list(intersect(rownames(full), cns17$V1)),
                            name = "cns17")
full <- AddModuleScore(full,        
                            features = list(intersect(rownames(full), cns18$V1)),
                            name = "cns18")
full <- AddModuleScore(full,        
                            features = list(intersect(rownames(full), cns19$V1)),
                            name = "cns19")
full <- AddModuleScore(full,        
                            features = list(intersect(rownames(full), cns20$V1)),
                            name = "cns20")

larvalcns$Tissue.Type <- factor(x = larvalcns$Tissue.Type, levels = c("collocytes",
                                                                      "PSCs",
                                                                      "PSCs related",
                                                                      "aATENs",
                                                                      "pATENs",
                                                                      "RTENs",
                                                                      "BTNs",
                                                                      "CESNs",
                                                                      "AMD+ motor ganglion",
                                                                      "Aristaless+ aSV",
                                                                      "Arx+ nerve cord (b)",
                                                                      "Arx+ pro-aSV",
                                                                      "Ci-hh2+ SV",
                                                                      "Ci-VP-R+ SV",
                                                                      "Ci-VP+ pSV",
                                                                      "coronet cells",
                                                                      "Dll-A+ neurohypophysis primordium",
                                                                      "eminence",
                                                                      "ependymal cells",
                                                                      "FoxD-b+ cells",
                                                                      "FoxP+ aSV",
                                                                      "GLGB+ pSV",
                                                                      "glia cells",
                                                                      "GLRA1/2/3+ motor ganglion",
                                                                      "GSTM1+ SV",
                                                                      "KCNB1+ motor ganglion",
                                                                      "Lhx1+ Bsh+ aSV",
                                                                      "Lhx1+ GABAergic neurons",
                                                                      "Lox5+ aSV",
                                                                      "MHB",
                                                                      "Opsin1+ PTPRB+ aSV",
                                                                      "Opsin1+ STUM+ aSV",
                                                                      "Pax2/5/8-A+ neck",
                                                                      "pigment cells",
                                                                      "Pitx+ neurohypophysis primordium",
                                                                      "Rx+ aSV",
                                                                      "Six3/6+ pro-aSV",
                                                                      "tail nerve cord (A)",
                                                                      "tail nerve cord (b)",
                                                                      "trunk nerve cord (A)",
                                                                      "trunk nerve cord (b)"))
dp <- DotPlot(larvalcns, features = c("cns01","cns21","cns31","cns41","cns51", "cns61","cns71","cns81","cns91","cns101","cns111","cns121","cns131","cns141","cns151","cns161","cns171","cns181","cns191", "cns201","pax37b01","pax37b11","pax37b21","pax37b31","pax37b41", "pax37b51","pax37b61","pax37b71","pax37b81","pax37b91"), group.by = "Tissue.Type")

ddf<- dp$data

### the matrix for the scaled expression 
dexp_mat <- ddf %>% 
  select(-pct.exp, -avg.exp) %>%  
  pivot_wider(names_from = id, values_from = avg.exp.scaled) %>% 
  as.data.frame() 

dexp_mat2 <- dexp_mat[,-1]
rownames(dexp_mat2) <- dexp_mat$features.plot

dexp <- as.matrix(sapply(dexp_mat2, as.numeric))  
rownames(dexp) <-  dexp_mat$features.plot


dexp <- as.matrix(dexp[,-1])
dexp <- na.omit(dexp)

rownames(dexp) <- str_sub(rownames(dexp), end=-2)

# plot the full dataset data
dpfull <- DotPlot(full, features = c("pax37b01","pax37b11","pax37b21","pax37b31","pax37b41", "pax37b51","pax37b61","pax37b71","pax37b81","pax37b91"), group.by = "cluster_tissue")

ddffull <- dpfull$data

### the matrix for the scaled expression 
dexp_matfull <- ddffull %>% 
  select(-pct.exp, -avg.exp) %>%  
  pivot_wider(names_from = id, values_from = avg.exp.scaled) %>% 
  as.data.frame() 

dexp_mat2full <- dexp_matfull[,-1]
rownames(dexp_mat2full) <- dexp_matfull$features.plot

dexpfull <- as.matrix(sapply(dexp_mat2full, as.numeric))  
rownames(dexpfull) <-  dexp_matfull$features.plot

dexpfull <- na.omit(dexpfull)
rownames(dexpfull) <- str_sub(rownames(dexpfull), end=-2)


# plot the full dataset data de genes
dpfull_pax <- DotPlot(full, features = c("X4_UP1", "X6_UP1", "X8_UP1", "X4_DOWN1","X6_DOWN1", "X8_DOWN1"), group.by = "cluster_tissue")

ddffull_pax <- dpfull_pax$data

### the matrix for the scaled expression 
dexp_matfull_pax <- ddffull_pax %>% 
  select(-pct.exp, -avg.exp) %>%  
  pivot_wider(names_from = id, values_from = avg.exp.scaled) %>% 
  as.data.frame() 

dexp_mat2full_pax <- dexp_matfull_pax[,-1]
rownames(dexp_mat2full_pax) <- dexp_matfull_pax$features.plot

dexpfull_pax <- as.matrix(sapply(dexp_mat2full_pax, as.numeric))  
rownames(dexpfull_pax) <-  dexp_matfull_pax$features.plot

dexpfull_pax <- na.omit(dexpfull_pax)
rownames(dexpfull_pax) <- str_sub(rownames(dexpfull_pax), end=-2)
rownames(dexpfull_pax) <- str_sub(rownames(dexpfull_pax), start=2)


dp_pax <- DotPlot(larvalcns, features = c("X4_UP1", "X6_UP1", "X8_UP1", "X4_DOWN1","X6_DOWN1", "X8_DOWN1"), group.by = "Tissue.Type")

ddf_pax<- dp_pax$data

### the matrix for the scaled expression 
dexp_mat_pax <- ddf_pax %>% 
  select(-pct.exp, -avg.exp) %>%  
  pivot_wider(names_from = id, values_from = avg.exp.scaled) %>% 
  as.data.frame() 

dexp_mat2_pax <- dexp_mat_pax[,-1]
rownames(dexp_mat2_pax) <- dexp_mat_pax$features.plot

dexp_pax <- as.matrix(sapply(dexp_mat2_pax, as.numeric))  
rownames(dexp_pax) <-  dexp_mat_pax$features.plot


dexp_pax <- as.matrix(dexp_pax[,-1])
dexp_pax <- na.omit(dexp_pax)

rownames(dexp_pax) <- str_sub(rownames(dexp_pax), end=-2)
rownames(dexp_pax) <- str_sub(rownames(dexp_pax), start = 2)

# make plots
fullplot <- pheatmap(t(dexpfull), border_color = "white", cellwidth = 10, cellheight = 10, cluster_rows = TRUE, cluster_cols = TRUE, treeheight_row = 6, treeheight_col = 6)
larvalcnsplot <- pheatmap(t(dexp), border_color = "white", cellwidth = 10, cellheight = 10, cluster_rows = TRUE, cluster_cols = TRUE, treeheight_row = 6, treeheight_col = 6)

full_pax <- pheatmap(t(dexpfull_pax), border_color = "white", cellwidth = 10, cellheight = 10, cluster_rows = TRUE, cluster_cols = FALSE, treeheight_row = 6, treeheight_col = 6)
larval_pax <- pheatmap(t(dexp_pax), border_color = "white", cellwidth = 10, cellheight = 10, cluster_rows = TRUE, cluster_cols = FALSE, treeheight_row = 6, treeheight_col = 6)

lay <- rbind(c(1,2,3,3),
             c(1,2,3,3),
             c(1,2,4,4),
             c(1,2,4,4))

pdf("./pax37b/plots/heatmap_scaled_expression_scores_oax37b_epiderm_cluster_markers_ciona.pdf", height = 16, width = 16)

grid.arrange(grobs=list(fullplot[[4]],
                        full_pax[[4]],
                        larvalcnsplot[[4]],
                        larval_pax[[4]]), layout_matrix = lay )
dev.off()
```


Finally to generate the clustered dotplot of Ciona robusta orthologs of adult Nasse markers for Fig. 11:


```
# set maximum ram usage:
options(future.globals.maxSize = 64000 * 1024^2)

# I set seed to ensure repeatability
set.seed(42)

library(circlize)
library(ComplexHeatmap)
library(data.table)
library(ggplot2)
library(patchwork)
library(plyr)
library(Polychrome)
library(reshape2)
library(Seurat)
library(tidyverse)
library(viridis)
library(clustree)


expr <- fread("/Users/davidlagman/Desktop/Integrated_data_seurat/ciona/expression_matrix_10stage.tsv", header = TRUE, row.names(1))
meta <- fread("/Users/davidlagman/Desktop/Integrated_data_seurat/ciona/edited_ciona10stage.cluster.upload.new.txt", header = TRUE)

rownames(meta) <- meta$NAME
rownames(expr) <- expr$GENE
expr <- as.data.frame(expr)
expr2 <- expr[,-1]
rownames(expr2) <- expr$GENE
full <- CreateSeuratObject(counts = expr2, meta.data = meta)


expr <- fread("/Users/davidlagman/Desktop/Integrated_data_seurat/ciona/CNS.lv.expressionmatrix.rename.tsv", header = TRUE, row.names(1))
meta <- fread("/Users/davidlagman/Desktop/Integrated_data_seurat/ciona/CNS.lv.clusters.upload.renameedit.txt", header = TRUE)

rownames(meta) <- meta$NAME
rownames(expr) <- expr$GENE
expr <- as.data.frame(expr)
expr2 <- expr[,-1]
rownames(expr2) <- expr$GENE
larvalcns <- CreateSeuratObject(counts = expr2, meta.data = meta)


orthnasse <- fread("/Users/davidlagman/Desktop/Integrated_data_seurat/final_results/nasse/tables/ciona_orthologs_sign.nasse.de.markers.txt", header = FALSE)

dp <- DotPlot(larvalcns, features = orthnasse$V1, group.by = "Tissue.Type")

ddf<- dp$data

### the matrix for the scaled expression 
dexp_mat <- ddf %>% 
  select(-pct.exp, -avg.exp) %>%  
  pivot_wider(names_from = id, values_from = avg.exp.scaled) %>% 
  as.data.frame() 

dpercent_mat<-ddf %>% 
  select(-avg.exp, -avg.exp.scaled) %>%  
  pivot_wider(names_from = id, values_from = pct.exp) %>% 
  as.data.frame() 

dexp_mat2 <- dexp_mat[,-1]
rownames(dexp_mat2) <- dexp_mat$features.plot
dpercent_mat2 <- dpercent_mat[,-1]
rownames(dpercent_mat2) <- dpercent_mat$features.plot

dexp <- as.matrix(sapply(dexp_mat2, as.numeric))  
rownames(dexp) <-  dexp_mat$features.plot
dperc <- as.matrix(sapply(dpercent_mat2, as.numeric))
rownames(dperc) <-  dpercent_mat$features.plot

## any value that is greater than 2 will be mapped to yellow
dcol_fun = circlize::colorRamp2(c(-2, 0, 2), viridis(20)[c(1,10, 20)])

dcell_fun = function(j, i, x, y, w, h, fill){
  grid.rect(x = x, y = y, width = w, height = h, 
            gp = gpar(col = NA, fill = NA))
  grid.circle(x=x,y=y,r= dperc[i, j]/100 * min(unit.c(w, h)),
              gp = gpar(fill = dcol_fun(dexp[i, j]), col = NA))}


mapnasse <- Heatmap(dexp,
                    heatmap_legend_param=list(title="avg expression", legend_direction = "vertical"),
                    column_title = "Larval CNS", 
                    col=dcol_fun,
                    rect_gp = gpar(type = "none"),
                    cell_fun = dcell_fun,
                    row_names_gp = gpar(fontsize = 5),
                    column_names_gp = gpar(fontsize = 5),
                    border = "black",
                    column_names_rot = 90)
mapnasse

fdp <- DotPlot(full, features = orthnasse$V1, group.by = "cluster_tissue")

fddf<- fdp$data

### the matrix for the scaled expression 
fdexp_mat <- fddf %>% 
  select(-pct.exp, -avg.exp) %>%  
  pivot_wider(names_from = id, values_from = avg.exp.scaled) %>% 
  as.data.frame() 

fdpercent_mat<-fddf %>% 
  select(-avg.exp, -avg.exp.scaled) %>%  
  pivot_wider(names_from = id, values_from = pct.exp) %>% 
  as.data.frame() 

fdexp_mat2 <- fdexp_mat[,-1]
rownames(fdexp_mat2) <- fdexp_mat$features.plot
fdpercent_mat2 <- fdpercent_mat[,-1]
rownames(fdpercent_mat2) <- fdpercent_mat$features.plot

fdexp <- as.matrix(sapply(fdexp_mat2, as.numeric))  
rownames(fdexp) <-  fdexp_mat$features.plot
fdperc <- as.matrix(sapply(fdpercent_mat2, as.numeric))
rownames(fdperc) <-  fdpercent_mat$features.plot

## any value that is greater than 2 will be mapped to yellow
fdcol_fun = circlize::colorRamp2(c(-2, 0, 2), viridis(20)[c(1,10, 20)])

fdcell_fun = function(j, i, x, y, w, h, fill){
  grid.rect(x = x, y = y, width = w, height = h, 
            gp = gpar(col = NA, fill = NA))
  grid.circle(x=x,y=y,r= fdperc[i, j]/100 * min(unit.c(w, h)),
              gp = gpar(fill = fdcol_fun(fdexp[i, j]), col = NA))}


fmapnasse <- Heatmap(fdexp,
                    heatmap_legend_param=list(title="avg expression", legend_direction = "vertical"),
                    column_title = "Full dataset", 
                    col=fdcol_fun,
                    rect_gp = gpar(type = "none"),
                    cell_fun = fdcell_fun,
                    row_names_gp = gpar(fontsize = 5),
                    column_names_gp = gpar(fontsize = 5),
                    border = "black",
                    column_names_rot = 90)
fmapnasse


pdf("/Users/davidlagman/Desktop/Integrated_data_seurat/final_results/plots/adult_nasse_markers_ciona_dataset.pdf", width = 20, height = 10)
grid.newpage()
pushViewport(viewport(layout = grid.layout(nr = 1, nc = 3)))
pushViewport(viewport(layout.pos.row = 1, layout.pos.col = 1:2))
draw(fmapnasse, newpage = FALSE)
upViewport()

pushViewport(viewport(layout.pos.row = 1, layout.pos.col = 3))
draw(mapnasse, newpage = FALSE)
upViewport()

pushViewport(viewport(layout.pos.row = 1, layout.pos.col = 3))
upViewport()

upViewport()
dev.off()
```
